# Polycomb-mediated repression of paternal chromosomes maintains haploid dosage in diploid embryos of Marchantia

**DOI:** 10.1101/2022.02.04.477531

**Authors:** Sean A. Montgomery, Tetsuya Hisanaga, Nan Wang, Elin Axelsson, Svetlana Akimcheva, Milos Sramek, Chang Liu, Frédéric Berger

## Abstract

Complex mechanisms regulate gene dosage throughout eukaryotic life cycles. Mechanisms controlling gene dosage have been extensively studied in animals, however it is unknown how generalizable these mechanisms are to diverse eukaryotes. Here, we use the haploid plant *Marchantia polymorpha* to assess gene dosage control in its short-lived diploid embryo. We show that throughout embryogenesis, paternal chromosomes are repressed resulting in functional haploidy. The paternal genome is targeted for genomic imprinting by the Polycomb mark H3K27me3 starting at fertilization, rendering the maternal genome in control of embryogenesis. Maintaining haploid gene dosage by this new form of imprinting is essential for embryonic development. Our findings illustrate how haploid-dominant species can regulate gene dosage through paternal chromosome inactivation and initiates the exploration of the link between life cycle history and gene dosage in a broader range of organisms.

## Introduction

Maintaining proper gene dosage is a challenge for eukaryotic organisms. For instance, multi-subunit protein complexes require balanced production of each component, lest incomplete, non-functional complexes are produced (Birchler & Veitia, 2010). Misregulation of gene dosage can lead to developmental defects, sterility, and disease (Loda, Collombet, & Heard, 2022). Dramatic changes in gene dosage notably occur during the process of diploidization after whole genome duplication (Edger & Pires, 2009) and sex chromosome evolution (Mank 2013). Sex chromosome dosage compensation is best understood mechanistically in mammalian female X chromosome inactivation (XCI) (Zylicz & Heard, 2020) and Drosophila male X chromosome upregulation (Samata & Akhtar, 2018). However, the molecular mechanisms are not conserved across the many diploid-dominant species in which sex chromosome dosage compensation has been described (Gu et al., 2019; Gu & Walters, 2017; Lau & Csankovszki, 2015; Lucchesi & Kuroda, 2015; Muyle et al., 2012) potentially due to the repeated innovation of sex chromosomes (Bachtrog et al., 2014). However, gene dosage also changes regularly during cell cycles and life cycles as ploidy levels change. Therefore, a large variety of gene dosage regulatory mechanisms remain to be discovered in eukaryotes.

All sexually reproducing eukaryotes have diploid and haploid life cycle stages, but the duration of each stage varies greatly amongst species. The alternation between ploidy must be programmed because unscheduled change in ploidy leads to genome instability (Davoli & de Lange, 2011). Despite the short haploid stage of gametes in mammals, gene dosage is managed by meiotic sex chromosome inactivation and post-meiotic silencing in male gametes (Lee & Bartolomei, 2013; Namekawa et al., 2006). This is continued as imprinted X chromosome inactivation (XCI) in early female embryos, wherein the male X chromosome is selectively repressed (Takagi & Sasaki, 1975). The disruption of meiotic sex chromosome inactivation results in meiotic arrest (Turner, 2007), illustrating its essentiality for sexual reproduction.

However, the mechanisms of gene dosage control throughout the mammalian life cycle are not conserved amongst animals (Maine, 2010; Turner, 2015; Vibranovski, 2014), reflective of the diversity of sex chromosome dosage compensation mechanisms (Gu et al., 2019; Gu & Walters, 2017; Lau & Csankovszki, 2015; Lucchesi & Kuroda, 2015; Muyle et al., 2012). Most eukaryotic life cycles differ from that of animals, with a predominance of haploid life stages, suggesting that there may be extensive diversity yet uncovered.

Haploid and haploid-dominant species present an intriguing and understudied corollary to understand gene dosage control throughout life cycles. Strictly haploid species such as yeast show limited evidence for gene dosage control (Chen et al., 2020; Hose et al., 2015; Springer, Weissman, & Kirschner, 2010). Haploid-dominant species with a short diploid phase of development are of particular interest because of the stark contrast of life cycles with diploid-dominant species and their prevalence across various branches of eukaryotic life. How, or even if, haploid-dominant species balance gene dosage during the diploid phase is not known.

Here, we uncovered a control of gene dosage by selective repression of alleles of paternal origin in the diploid embryonic stage of the model haploid-dominant bryophyte *Marchantia polymorpha* (hereafter referred to as Marchantia). We show that Marchantia represses paternal chromosomes by genomic imprinting via the Polycomb mark H3K27me3, the first description of imprinting in the bryophyte lineage since its theoretical prediction (Carey, Kollar, & McDaniel, 2021; Haig, 2013; Haig & Wilczek, 2006; Montgomery & Berger, 2021; Shaw, Szovenyi, & Shaw, 2011). Disruption of this unique form of genomic imprinting, which we term “paternal chromosome inactivation” (PCI), results in derepression of the paternal genome and lethality. Furthermore, we show that the imprinting mark is deposited at the pronuclear stage and initiates PCI that persists until the end of embryogenesis. Therefore, Marchantia manages gene dosage by effectively maintaining a functionally haploid state in diploid embryos under the control of the maternal genome.

## Results

### Embryonic transcription is maternally biased

To explore Marchantia gene dosage control, we performed crosses between two wild-type natural accessions, Cam-2 as the mother and Tak-1 as the father, and obtained transcriptomes from embryos thirteen days after fertilization (daf) (Figure 1A). Each transcriptome was prepared from single hand-dissected embryos that were washed several times to remove potential contaminating RNA from the surrounding maternal tissue (Figure 1B; Video S1) (Schon & Nodine, 2017). A comparison of embryonic, vegetative, and sexual organ transcriptomes demonstrated the distinctness of embryonic transcriptomes from other tissues and the similarity of each embryonic transcriptome to each other (Figures 1C and Figure 1-figure supplement 1A). Together, these results indicated to us that we had obtained pure embryonic transcriptomes for further analyses.

**Figure 1.**
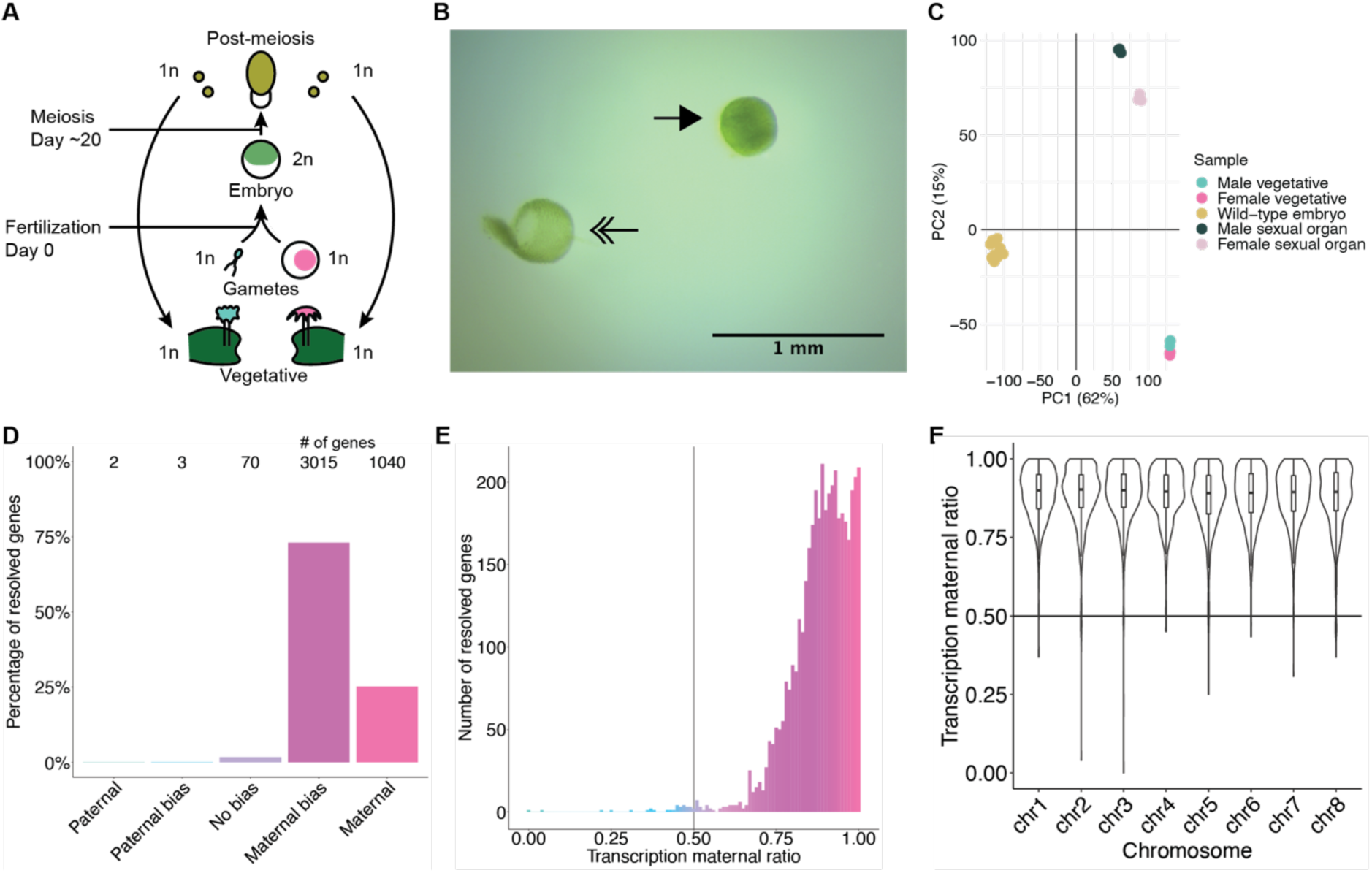
Embryonic transcription is maternally biased. (**A**) Life cycle of *Marchantia polymorpha*. Haploid (1n) vegetative males and females produce male and female reproductive structures, which subsequently produce sperm and egg. The diploid (2n) embryo persists for around 20 days before meiosis and the production of haploid spores. Ploidy of each stage is indicated. (**B**) Image of a representative hand-dissected embryo after removal of perianth and calyptra of maternal origin. Solid single arrow indicates isolated embryo. Double arrow indicates the removed calyptra. Scale bar as indicated. (**C**) Principal component analysis of transcriptomes from wild-type embryos (Cam-2 x Tak-1), vegetative tissues from female and male parents, and female and male sexual organs. The first two principal components are plotted, and the percentage of variance explained is indicated. (**D**) Percentage of measured genes within each category of maternal ratio (*pm*) of transcription in wild-type embryos. Segments are for paternal (*pm* < 0.05), paternal bias (0.05 < *pm* ≤ 0.35), no bias (0.35 < *pm* < 0.65), maternal bias (0.65 ≤ *pm* < 0.95), and maternal (0.95 ≤ *pm*) expression of genes, with the number of genes indicated above each bar. (**E**) Histogram of the maternal ratio (*pm*) of transcription per gene in wild-type (Cam-2 x Tak-1) embryos. Each bin is 0.01 units wide. (**F**) Violin plots of transcription maternal ratio of genes per chromosome. Sex chromosomes are excluded as alleles could not be resolved. See also Figure 1-figure supplement 1

We further looked for evidence of allele-specific expression in diploid embryos. We utilized single-nucleotide polymorphisms (SNPs) between female and male accessions to calculate the ratio of reads originating from maternal alleles versus paternal alleles (maternal ratio, *p_m_*; ascribed a value between 0 and 1, ranging from 0 if only paternal reads were detected to 1 if only maternal reads were detected). Combining all replicates, we only considered genes with at least fifty reads containing informative SNPs. Transcription was overall maternally biased for 98% of resolved genes, with 73% of genes maternally biased (as defined in (X. Wang & Clark, 2014); 0.65 ≤ *p_m_* < 0.95) and 25% of genes only expressed from maternal alleles (*p_m_* ≥ 0.95) (Figures 1D-E). The strong unidirectional bias in gene expression suggested homogeneity amongst replicates, which was confirmed when assessing the maternal ratio of transcription from each replicate (Figure 1-figure supplement 1B). We conclude that in Marchantia embryos, genes are primarily or exclusively expressed from their maternal allele.

The exact reciprocal cross was not possible because inbred genetically near-identical pairs of males and females do not exist, but to confirm that the maternal bias did not result from the pair of natural accessions used, we analyzed published RNA-seq data from a cross of different accessions, Tak-2 and Tak-1 (Frank & Scanlon, 2015). The published transcriptomes were generated from samples collected by laser-capture microdissection, an orthogonal sample collection method that offered equally high sample purity (Schon & Nodine, 2017). A similarly strong maternal bias in transcription was observed, with 99% of genes maternally biased or expressed only from maternal alleles (Figure 1-figure supplements 1C-D). Thus, these data ruled out that the observed allele-specific gene expression originated from natural variation amongst wild-type parents or from maternal contamination during sample collection. Additionally, we tested whether the genes resolved by our analyses formed a sample representative of all genes. We found no correlation between the maternal ratio and expression level of a gene (Figure 1-figure supplement 1E), nor did the transcription maternal ratio vary significantly along the length of each autosome (Figure 1F and Figure 1-figure supplement 1F). Thus, we infer that the genes we were able to resolve with SNPs were representative of a genome-wide maternal bias in transcription. Overall, the lack of paternal allele expression suggests the presence of a repressive chromatin modification specifically on the paternal genome.

### Levels of H3K27me3 enrichment are paternally biased

To better understand what chromatin-related mechanisms may be driving the maternal bias in embryonic transcription, we examined differentially expressed genes between vegetative parents and embryos. In total, 3879 genes were upregulated in embryos relative to both mothers and fathers (Figure 2-figure supplement 1A), while 3466 genes were downregulated (Figure 2-figure supplement 1B). Upregulated genes were more expressed from the maternal genome than downregulated genes (Figure 2-figure supplement 1C, effect size (Cohen’s *d*) = 0.276) highlighting a maternal control over the embryonic transcriptome. Since imprinting is an epigenetic process, we focused further on chromatin-related genes. Of the 215 chromatin-related genes in the Marchantia genome (Bowman et al., 2017), 151 were upregulated and 7 were downregulated (Figure 2-figure supplement 1D; Table S1). Of these, 20 genes were specifically expressed in the embryonic stage (Table S1, Transcripts per Million greater than 1 in embryos and less than 1 in other tissues). Two noteworthy genes were paralogs of the catalytic subunit of the Polycomb Repressive Complex 2 (PRC2), *E(z)2* and *E(z)3* (Figure 2A). PRC2 is a conserved multi-subunit complex that deposits H3K27me3 and is associated with gene silencing (Margueron & Reinberg, 2011). The other three subunits of PRC2, *FIE*, *Su(z)12*, and *MSI1*, and a third catalytic subunit paralog, *E(z)1*, were expressed in all tissues (Figure 2A). Therefore, we hypothesized that H3K27me3 might be present on silenced paternal alleles in the Marchantia embryo.

**Figure 2.**
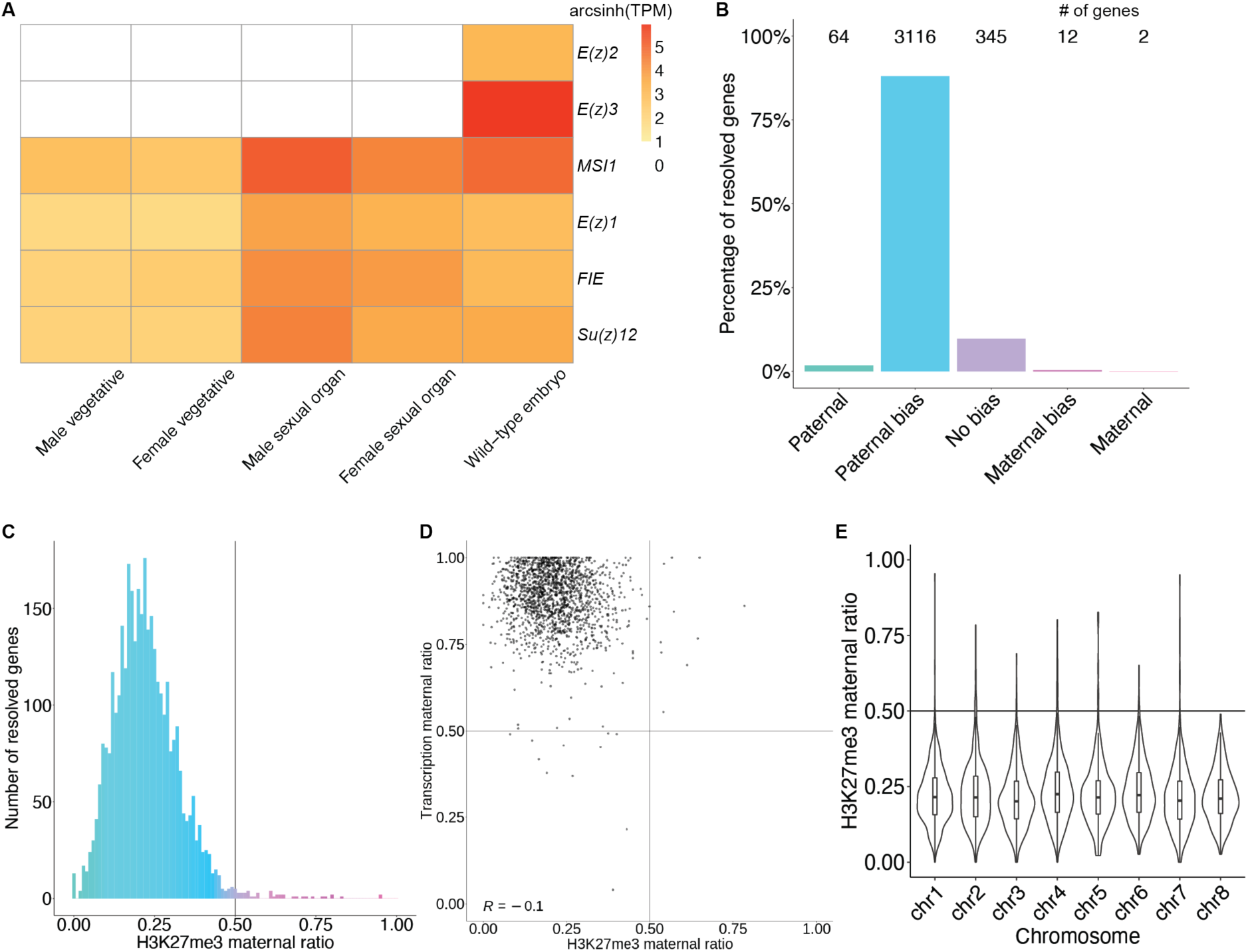
Levels of H3K27me3 enrichment are paternally biased. (**A**) Heatmap of gene expression of Polycomb Repressive Complex 2 subunits across Marchantia development. Vegetative male (Tak-1) and female (Cam-2) tissues give rise to male and female sexual organs (antheridiophores and archegoniophores, respectively; data from (Higo et al., 2016)). Wild-type embryos are from Cam-2 x Tak-1 crosses. Values shown are arcsinh transformed Transcript per Million values. (**B**) Percentage of measured genes within each category of maternal ratio (*p_m_*) of H3K27me3 in wild-type embryos. Segments are for paternal (*p_m_* < 0.05), paternal bias (0.05 < *p_m_* ≤ 0.35), no bias (0.35 < *p_m_* < 0.65), maternal bias (0.65 ≤ *p_m_* < 0.95), and maternal (0.95 ≤ *p_m_*) H3K27me3 of genes, with the number of genes indicated above each bar. (**C**) Histogram of the maternal ratio (*p_m_*) of H3K27me3 per gene in wild-type (Cam-2 x Tak-1) embryos. Each bin is 0.01 units wide. (**D**) Scatterplot of maternal ratios of H3K27me3 and transcription per resolved gene. Spearman correlation is indicated. (**E**) Violin plots of H3K27me3 maternal ratio of genes per chromosome in wild-type embryos. Sex chromosomes are excluded as alleles could not be resolved. See also Figure 2-figure supplement 1

We first set out to determine whether H3K27me3 was enriched on paternal alleles in Marchantia embryos. We profiled chromatin modifications using CUT&RUN (Skene & Henikoff, 2016; Zheng & Gehring, 2019) on sorted nuclei from Marchantia embryos. We used SNPs between male and female accessions to distinguish the parental allele of origin for CUT&RUN reads to calculate a maternal ratio (*p_m_*) for genomic regions of interest. Enrichment in H3K27me3 was paternally biased for 88% of genes resolved (0.05 < *p_m_* ≤ 0.35) (Figure 2B-C) and genes with paternally biased H3K27me3 had maternally biased transcription (Figure 2D). Genes paternally marked with H3K27me3 were located across all autosomes (Figure 2E and Figure 2-figure supplement 1E), indicating the broad, pervasive nature of the phenomenon. In contrast, a paternal bias was not observed in profiles of H3K9me1, H3, and H3K36me3 (5%, 2%, and 2% of genes with 0.05 < *p_m_* ≤ 0.35, respectively) (Figure 2-figure supplement 1F-H). We conclude that levels of H3K27me3 enrichment anticorrelate with maternally biased transcription and spreads over most paternal alleles in Marchantia embryos. These findings suggest that H3K27me3 covers the entire genome of paternal origin.

### Partitioning of the paternal genome into dense H3K27me3 compartments

To test if the paternal genome was coated with H3K27me3, we performed immunofluorescence experiments to observe the localization of this modification within embryonic nuclei. As a control, nuclei of parental adult vegetative cells showed evenly distributed speckles of heterochromatin marked by H3K27me3 (Figure 3A) (Montgomery et al., 2020). In stark contrast, one to three large heterochromatic compartments, as defined by dense DNA staining, covered 10% of the area in embryonic nuclei (Figure 3A and Figure 3-figure supplement 1A-B). A strong correlation between heterochromatic foci and H3K27me3 was apparent (Figure 3A). 44% of the H3K27me3 signal was contained within these compartments (Figure 3-figure supplement 1C), whereas 70% of the area of the compartments was contained within H3K27me3 domains (Figure 3-figure supplement 1D). In contrast, only 20% and 10% of H3K9me1 and H3K36me3 signal, respectively, were contained within heterochromatic compartments (Figure 3-figure supplement 1C). H3K9me1 is indicative of constitutive heterochromatin on repetitive genomic regions in Marchantia and other eukaryotes, whereas H3K36me3 is associated with expressed genes (Montgomery et al., 2020). We conclude that the portion of the genome marked by H3K27me3 represents the largest fraction of heterochromatin organized as a couple of dense compartments in embryonic nuclei.

**Figure 3.**
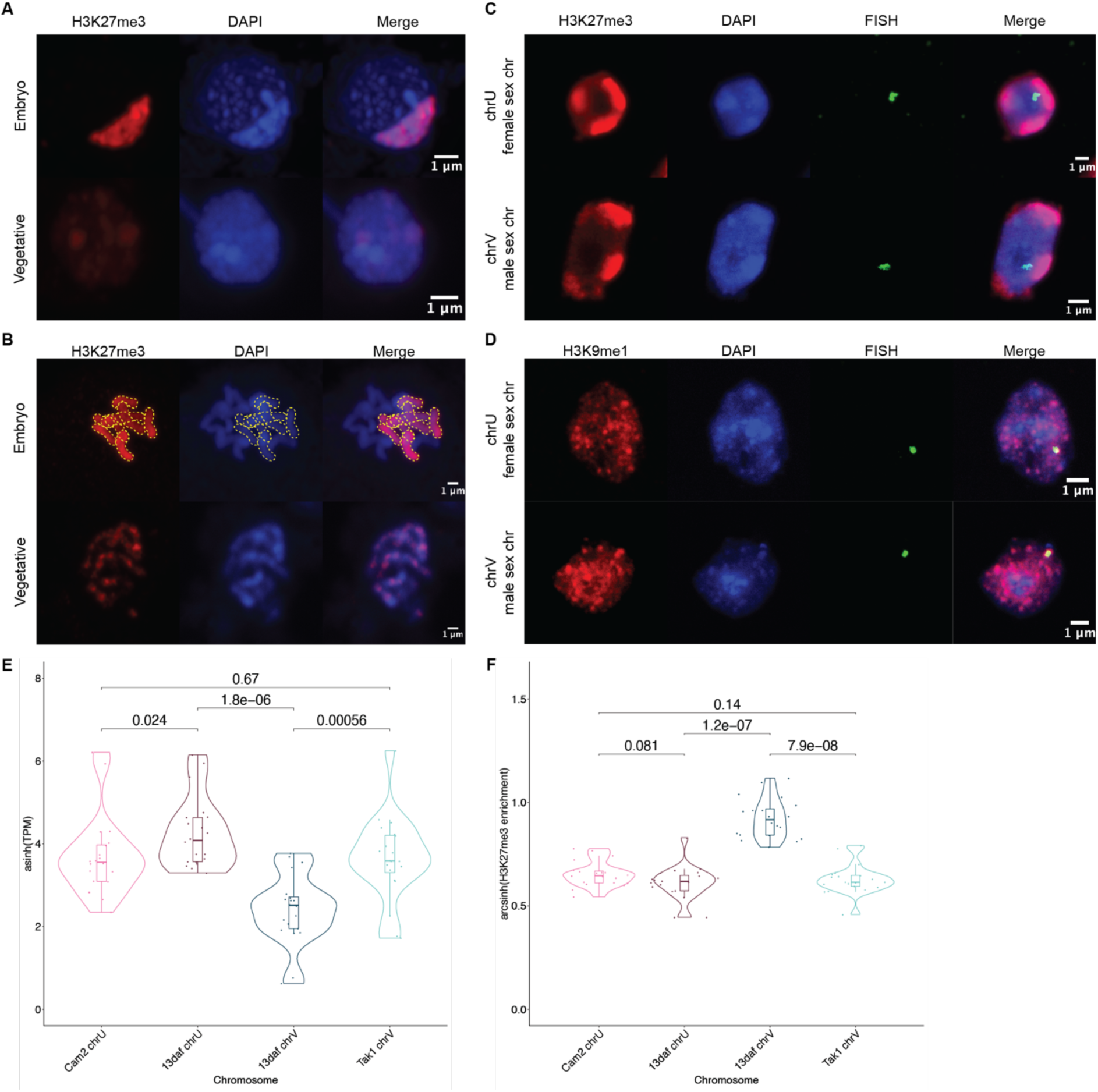
Paternal autosomes are coated in H3K27me3 and partitioned in heterochromatic foci. (**A**) Immunofluorescence of H3K27me3 in interphase wild-type embryonic and vegetative nuclei. DNA is stained with DAPI. Scale bar as indicated. (**B**) Immunofluorescence of H3K27me3 in mitotic wild-type embryonic and vegetative cells. DNA is stained with DAPI. Contrast was enhanced for the DAPI channel of vegetative nuclei for illustration purposes. Outlines of the H3K27me3-coated chromosomes are indicated with dashed yellow lines. Scale bar as indicated. (**C**) Immuno-FISH for sex chromosomes and H3K27me3 in interphase wild-type embryonic nuclei. The female sex chromosome is chrU and the male sex chromosome is chrV. Scale bar as indicated. (**D**) Immuno-FISH for sex chromosomes and H3K9me1 in interphase wild-type embryonic nuclei. The female sex chromosome is chrU and the male sex chromosome is chrV. Scale bar as indicated. (**E**) Violin plot of arcsinh transformed Transcript per Million (TPM) values for sex chromosome gametologs in vegetative (Cam2 and Tak1) and embryonic (13 days after fertilization (daf)) samples. *P* values are indicated, unpaired two-tailed Wilcoxon test. (**F**) Violin plot of arcsinh transformed H3K27me3 enrichment for sex chromosome gametologs in vegetative and embryonic samples. *P* values are indicated, unpaired two-tailed Wilcoxon test. See also Figure 3-figure supplement 1

### All paternal autosomes are coated with H3K27me3

The Marchantia life cycle has a vegetative haploid phase with a diploid phase of embryogenesis (Figure 1A). In Marchantia diploid embryonic cells, the genome is packaged in 2n = 18 chromosomes including two sex chromosomes and sixteen autosomes. The large size of the heterochromatic compartments marked by H3K27me3 suggested that they contained entire chromosomes. Accordingly, immunofluorescence of mitotic cells revealed that eight of the sixteen autosomes were densely coated with H3K27me3, whereas the other half had no detectable H3K27me3 (Figure 3B). In contrast, mitotic vegetative haploid cells showed an uneven speckled pattern of H3K27me3 over the eight autosomes (Figure 3B). This observation mirrored the strong paternal bias of H3K27me3 enrichment, and we conclude that the paternal genome is covered by H3K27me3 and partitioned into heterochromatic compartments within embryonic nuclei.

In addition to the eight autosomes, each parent carries a small sex chromosome (U in females and V in males (Iwasaki et al., 2021)). Sex chromosomes detected using FISH were not associated with H3K27me3 heterochromatic foci in Marchantia embryos (Figure 3C and Figure 3-figure supplement 1E). Instead, we observed that both U and V sex chromosomes rather associated with H3K9me1 heterochromatic foci (Figure 3D and Figure 3-figure supplement 1E). We conclude that the sex chromosomes are excluded from H3K27me3 heterochromatic compartments and form small constitutive heterochromatic foci in both embryonic and vegetative nuclei. Yet, the protein coding genes on the female U sex chromosome are expressed at a much higher level than homologous genes on the male V sex chromosome (Figure 3E). This imbalance towards female expression is correlated with an enrichment of H3K27me3 on the genes of the male chromosome (Figure 3F and Figure 3-figure supplement 1F). Hence, overall, H3K27me3 targets the paternal alleles of all chromosomes in Marchantia, resulting in a pseudo-haploid state in the embryo.

### H3K27me3 is deposited in paternal pronuclei

Like in most animals, the male pronucleus contributed by the sperm remains separated from the female pronucleus contributed by the egg in the Marchantia zygote, thus providing an opportunity for the deposition of an epigenetic mark on the genome of one parent (Hisanaga et al., 2021, 2019). As Marchantia sperm chromatin is comprised of protamines and is devoid of histones (D’Ippolito et al., 2019; Reynolds & Wolfe, 1978), we hypothesized that paternal H3K27me3 is deposited on paternal alleles sometime after fertilization. Male and female pronuclei remain separate until 4 daf (Figure 4A-B) (Hisanaga et al., 2021), thus we examined if H3K27me3 was deposited at 3 daf, before pronuclear fusion. As we could not isolate pronuclei for chromatin profiling, we instead performed immunofluorescence experiments. At 3 daf, we observed both H3K27me3 and H3 in the paternal pronucleus (Figure 4C and Figure 4-figure supplement 1), demonstrating that H3K27me3 is deposited in the paternal pronucleus before its fusion with the maternal pronucleus. Therefore, paternal alleles become imprinted by H3K27me3 while they are spatially segregated from maternal alleles prior to the fusion of pronuclei. The conservative restoration of H3K27me3 after DNA replication (Jiang & Berger, 2017) provides a mechanism to propagate the initial paternal “coat” of H3K27me3 to all autosomes and silence the paternal genome throughout embryonic development.

**Figure 4.**
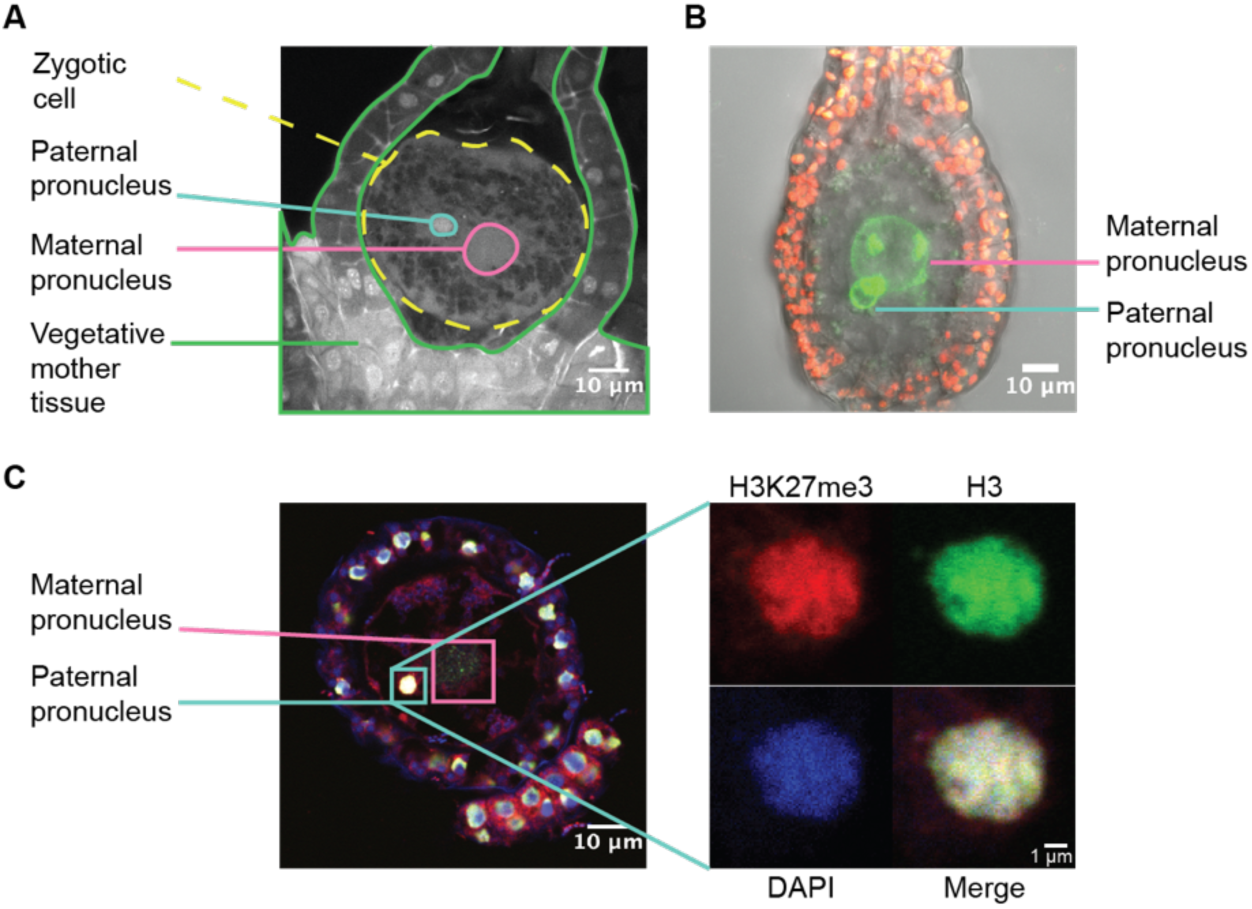
H3K27me3 is deposited in paternal pronuclei. (**A**) Annotated confocal image of a Marchantia zygote 3 days after fertilization (daf) with surrounding vegetative mother tissue. The paternal pronucleus is visible in the vicinity of the maternal pronucleus. Nuclei are stained with DAPI. Indicated are the fertilized zygotic cell (dashed yellow circle), maternal pronucleus (pink circle), vegetative mother tissue (green lines) surrounding the zygote, and paternal pronucleus (cyan circle). Scale bar as indicated. (**B**) Composite maximum intensity projection confocal image of a Marchantia zygote expressing *SUN-GFP* at 3 daf plus surrounding vegetative mother tissue. Nuclear membranes are marked by localization of SUN-GFP, shown in green. The paternal pronucleus is smaller than and adjacent to the maternal pronucleus. Autofluorescence from chloroplasts in vegetative mother cells is shown in red, and both channels are overlayed on a transmitted light image. Scale bar as indicated. (**C**) Immunofluorescence image 3 daf of a Marchantia zygote. Both maternal and paternal pronuclei are indicated in pink and cyan, respectively. The inset depicts a zoomed in view of the paternal pronucleus with separate images for H3K27me3 (red), H3 (green), DAPI (blue), and the merged image. Contrast is enhanced for each image and channel independently for visualization purposes. Scale bars as indicated. See also Figure 4-figure supplement 1

### Embryonic PRC2 deposits H3K27me3 and represses the paternal genome

To directly test the effect of embryo-specific PRC2 subunits on paternal H3K27me3 imprinting, we knocked-out both embryo-specific paralogs of the catalytic subunit, *E(z)2* and *E(z)3* (Figure 5-figure supplement 1A). These mutants did not display any aberrant phenotype prior to fertilization (Figure 5-figure supplement 1B-C). We crossed mutant females (Cam-2 *e(z)2/e(z)3*) to wild type males and observed a loss or dispersion of large heterochromatic foci that overlapped with H3K27me3 foci in embryos compared to wild-type embryos (Figure 5A and Figure 5-figure supplement 1D). In these embryos, enrichment of H3K27me3 as measured by CUT&RUN was significantly decreased over most genes (Figure 5-figure supplement 1E, Wilcoxon signed-rank test p < 0.0001). Thus, maternal inheritance of both *e(z)2* and *e(z)3* altered patterns of H3K27me3 associated heterochromatic foci and reduced H3K27me3 deposition. The paralog *E(z)1* is expressed in embryos (Figure 2A) and likely accounted for the remaining detected H3K27me3, but it proved impossible to test this hypothesis due to lethality at the haploid vegetative stage in knockdowns of *E(z)1* (Flores-Sandoval, Dierschke, Fisher, & Bowman, 2016). To determine whether paternal alleles were the source of H3K27me3 loss, we distinguished the parental genome of origin of CUT&RUN sequencing reads and calculated maternal ratios. The paternally biased enrichment of H3K27me3 was significantly reduced (Figure 5B, Wilcoxon signed-rank test p < 0.0001, effect size = 1.03) and only 51% of genes were categorized as paternally biased (0.05 < *p_m_* ≤ 0.35) (Figure 5-figure supplement 1F), down from 88% in wild type (Figure 2B). Furthermore, there was a negative correlation between H3K27me3 enrichment and maternal ratio (Figure 5C), indicating that loci that lost H3K27me3 had predominantly lost paternal H3K27me3 in embryos that lost the maternal alleles of *E(z)2* and *E(z)3*. We conclude that the deposition of H3K27me3 on most paternal loci in Marchantia embryos depends on maternally supplied PRC2 activity.

**Figure 5.**
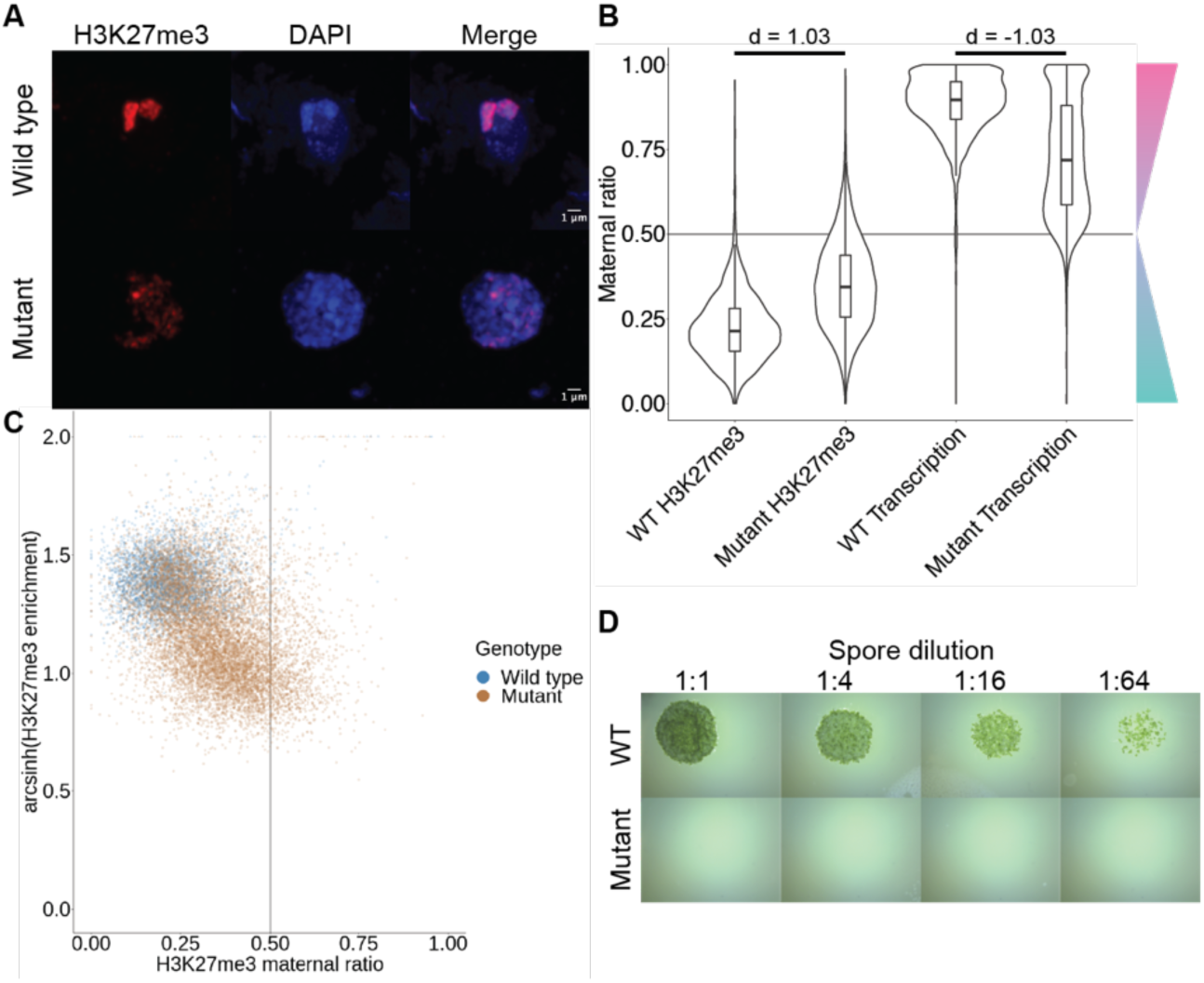
Embryonic PRC2 deposits H3K27me3 and represses the paternal genome. (**A**) Immunofluorescence of H3K27me3 in interphase wild-type and mutant embryonic nuclei. DNA is stained with DAPI. (**B**) Violin plots of maternal ratios for wild-type (WT) and mutant H3K27me3 and transcription. Cohen’s *d* effect size values are indicated for pairwise comparisons of mutant to WT H3K27me3 maternal ratio and mutant to WT transcription maternal ratio, where |d| > 0.8 is a large effect, as previously reported (Cohen, 1992). (**C**) Scatterplot of H3K27me3 enrichment versus H3K27me3 maternal ratio per gene in WT and mutant embryos. Genes with an arcsinh-transformed H3K27me3 enrichment greater than 2 are displayed as triangles at the upper boundary of the plot. (**D**) Spore germination assay for spores resulting from WT and mutant embryos. A serial dilution of a suspension of spores from a single embryo is shown. See also Figure 5-figure supplements 1, 2

If H3K27me3 did indeed repress paternal alleles, we expected expression from paternal alleles at loci that lost paternal H3K27me3 in *e(z)2/e(z)3* mutants. To test this idea, we generated transcriptomes from mutant embryos and examined maternal ratios of gene expression. Overall, transcription became more biallelic in mutants than in wild-type embryos (Figure 5B, Wilcoxon signed-rank test p < 0.0001, effect size = -1.03). Only 47% of genes were maternally biased (0.65 ≤ *p_m_* <0.95) and 15% of genes completely expressed from maternal alleles (*p_m_* ≥ 0.95), a stark deviation from wild type values of 73% and 25% (Figure 5-figure supplement 2A and compare with Figure 1D). Comparing both mutant maternal ratios of transcription with patterns of H3K27me3, we observed that the H3K27me3 maternal ratio negatively correlated with the maternal ratio of transcription (Figure 5-figure supplement 2B-C) and that H3K27me3 enrichment positively correlated with the maternal ratio of transcription (Figure 5-figure supplement 2B, D), indicating that loci with less paternal H3K27me3 in mutants were more transcribed from paternal alleles. High paternal H3K27me3 and maternal transcription did not correlate with the level of gene expression (Figure 5-figure supplement 2B, E-G), suggesting that absolute gene expression levels did not influence paternal allele repression. Therefore, paternal alleles regain expression in the absence of maternal PRC2 and upon the loss of paternal H3K27me3. We conclude that H3K27me3 deposited by embryo-specific PRC2 subunits is required for the collective repression of paternal alleles, resulting in paternal chromosome inactivation (PCI).

To assess the physiological relevance of the loss of paternal allele repression, we quantified the growth and survival of mutant embryos. Embryonic growth was significantly slower in mutants than in wild type, as measured by total size (Figure 5-figure supplement 2H). Only 20% of mutant embryos survived to maturity versus 95% of wild type (Figure 5-figure supplement 2I), and only 5% of all mutant embryos produced spore-bearing structures, compared to 77% of wild type (Figure 5-figure supplement 2J). Of those mutants that produced spores, none produced viable spores, thus rendering all mutants unable to continue their life cycle (Figure 5D). We conclude that our results are consistent with the model that PRC2-mediated PCI is essential for viability and fecundity in Marchantia (Figure 6).

**Figure 6.**
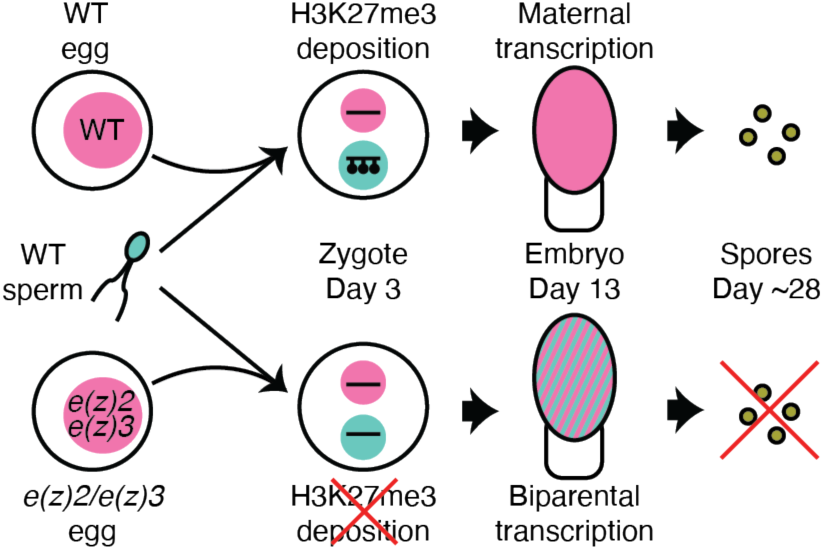
Model of genomic imprinting in Marchantia. Model of H3K27me3 deposition in WT paternal pronuclei and subsequent propagation throughout embryogenesis. Closed lollipops depict H3K27me3 on genes. Pink and blue circles depict maternal and paternal (pro)nuclei, respectively. Pink and striped ovals depict whole embryos and the parental genome from which transcription is occurring. Yellow circles depict mature spores. The lack of H3K27me3 on paternal pronuclei in mutant zygotes allows for the transcription of paternal alleles in the embryo, ultimately leading to the lack of viable spore production.

## Discussion

In the present study, we report how a haploid-dominant species controls gene dosage during a short diploid stage. While diploid-dominant species often balance gene dosage between autosomes and sex chromosomes throughout their life cycles, haploid-dominant species must manage gene dosage only during embryonic development. The model bryophyte *Marchantia polymorpha* achieves gene dosage control during embryonic development via genomic imprinting and subsequent repression of the paternal genome. The repressive mark H3K27me3 is first deposited over the whole paternal pronucleus, in contrast to mammals where H3K27me3-mediated imprinting impacts only a handful of loci and is deposited in the female gamete (Inoue, Jiang, Lu, Suzuki, & Zhang, 2017). Interference of maternal PRC2-mediated H3K27me3 deposition ultimately halts embryonic development. Thus, PRC2 initiates and maintains silencing of the entire paternal genome, which is essential for the development of the diploid embryo of Marchantia.

The bryophyte life cycle marks the transition between the haploid life cycle of their aquatic ancestors and the diploid life cycle of vascular plants. Our results suggest that maintaining the dominant haploid dosage of gene expression was selected in the diploid embryo of bryophytes. In response to whole genome duplications, plant and animal genomes modulate gene expression to pre-duplication levels (McElroy et al., 2017; Pala, Coelho, & Schartl, 2008; Song, Potter, Doyle, & Coate, 2020), though the mechanisms underlying such gene dosage control are poorly understood. Why gene dosage control in Marchantia was achieved through imprinting might be explained by the fact that ancestor of bryophytes had separate sexes (Iwasaki et al., 2021) and embryos developed on mothers. It is thus likely that viviparity and its collateral maternal support of embryo development favored the evolution of imprinting as a way to impose maternal control, as proposed by a body of theoretical works (Carey et al., 2021; Haig, 2013; Haig & Wilczek, 2006; Montgomery & Berger, 2021; Shaw et al., 2011). Imprinting has not been discovered in viviparous non-therian animals, but only orthologous loci imprinted in mammals were investigated, thus a genome-wide search may yield new insights (Griffith, Brandley, Belov, & Thompson, 2016; Lawton et al., 2005; Renfree, Suzuki, & Kaneko-Ishino, 2013). Our findings support the idea that viviparity is sufficient for the evolution of imprinting and provide a new framework to explore the evolution of imprinting in a much more diverse range of organisms than previously considered across eukaryotes.

We believe that PCI represents a new form of imprinting. It is distinct from parental genomic imprinting described in flowering plants and therian mammals for three main reasons: the epigenetic mark is deposited after fertilization; the entire genome of one parent is silenced by the epigenetic mark; and imprinting imposes a global maternal control of embryogenesis. The outcome of imprinting in Marchantia also differs from the elimination of the paternal genome following heterochromatin formation in several insect species (Crouse, 1960; de la Filia et al., 2021). In insects, heterochromatinized paternal chromosomes are eliminated by still unknown mechanisms, though the timing of elimination varies amongst species leading to pseudohaploidy in some cases (Bain et al., 2021; Morse & Normark, 2005). In contrast, the paternal genome in Marchantia embryos subjected to PCI is reactivated and still passed on to the next generation.

PCI in Marchantia differs from XCI in mammals because all paternal chromosomes are compacted and repressed and H3K9me is not involved (Heard et al., 2001). The formation of heterochromatic foci associated with H3K27me3 is reminiscent of the compaction and compartmentalization of the X chromosome during XCI in mammals (Galupa & Heard, 2018; Nozawa et al., 2013; Plath et al., 2003). However, since half of the genome is marked exclusively by H3K27me3, its partition results in large compartments, the compaction of which depends on PRC2 activity. Similar mechanisms may be at play in mediating whole-chromosome compaction and repression in mammals, insects and Marchantia. The precise molecular mechanisms underlying the establishment and abolition of PCI are not addressed by our model (Figure 6), however their elucidation will be of interest to make cross-kingdom comparisons with other instances of imprinted dosage compensation mechanisms.

Overall, we have uncovered a distinct mechanism that controls gene dosage in a haploid-dominant species. We anticipate similar controls for all bryophytes as well as other groups of organisms that alternate long lived haploid and diploid phases. Remnants of such a control might exist in flowering plants, as suggested by maternally dominant expression in the rice zygote (Anderson et al., 2017). Yet, both parental genomes in Arabidopsis are equally expressed after fertilization (Schon & Nodine, 2017), but whether the total dosage of expression is the same as in haploid progenitors of gametes remains unknown. Various forms of alternation between multicellular haploid and diploid life phases are also widespread in brown and red algae. Brown algae show changes in epigenetic marks and transcription between haploid and diploid generations, despite the absence of Polycomb (Bourdareau et al., 2021). Thus, it would be of interest to determine the mechanisms of gene dosage control in these species as it would be distinct from PCI in Marchantia. Broadly, we propose that although sex chromosomes provide an important paradigm to understand gene dosage control, this phenomenon evolved several times as life cycles alternating between ploidy levels diversified, suggesting that there is an expanse of gene dosage regulatory mechanisms that remains to be explored across the broad assortment of eukaryotic life cycles.

## Materials and Methods

### Plant lines and growth conditions

Wild-type male Tak-1, female Cam-2, and female Tak-2 accessions of *Marchantia polymorpha* ssp. *ruderalis* were used in this study. Cam-2 and Tak-2 *e(z)2/e(z)3* mutants were generated as described below.

Female plants for crosses were grown at room temperature on Grodan Vital (Grodan, Roermond, The Netherlands) supplemented with liquid Hyponex fertilizer (Hyponex, Osaka, Japan) under constant white light. Male plants for crosses were grown at 22 °C on Neuhaus N3 substrate soil (Humko, Podnart, Slovenia) under 16 hours of far-red light and 80% humidity. Plants grown for the collection or observation of vegetative tissues were grown under axenic conditions on half-strength Gamborg B5 media without vitamins (Duchefa Biochemie, Haarlem, The Netherlands) and 1% (w/v) agar under constant white light.

Crosses were performed by collecting mature antheridiophore discs in water and pipetting the water containing released sperm onto archegoniophores.

### Generation of e(z)2/e(z)3 mutants

To construct a plasmid to disrupt *E(z)2* and *E(z)3* simultaneously, two oligonucleotide pairs (TH219: ctcgAAATAGAAAGTGGCGCCT/TH220: aaacAGGCGCCACTTTCTATTT for *E(z)2*; TH223: ctcgATCATATACCCTCGGCTC /TH224:

aaacGAGCCGAGGGTATATGAT for *E(z)3*) were annealed and cloned into the BsaI sites of pMpGE_En04 and pBC-GE14 to yield pMpGE_En04-MpEz2-sg1 and pBC-GE14-MpEz3-sg1, respectively. These two plasmids were assembled via BglI restriction sites and ligated to yield pMpGE_En04-MpEz2-sg1-MpEz3-sg1. The resulting DNA fragment containing two MpU6promoter-gRNA cassettes was transferred into pMpGE010 (cat. no. 71536, Addgene) (Sugano et al., 2018) using the Gateway LR reaction (Thermo Fisher Scientific Inc, Waltham, MA, USA) to yield pMpGE010_MpEz2-sg1-MpEz3-sg1. This construct was introduced into Cam-2 gemmae using the G-AgarTrap method (Tsuboyama, Nonaka, Ezura, & Kodama, 2018). Transformants were selected for on 0.5 Gamborg B5 plates without vitamins (Duchefa Biochemie) supplemented with hygromycin and genotyped using the following primer pairs: TH300: TACGCCCTCTCCCATTGAAC/TH301: GATACGAAGAGAACGAACCTGC for *E(z)2* and TH306: TGAGCTACATGGCTACTCTCAACC/TH307: AGCTTGGAACACGGATCTCCTG for *E(z)3*.

### Transcriptome generation

Vegetative samples from Cam-2 and Tak-1 were collected from 100mg of apical notches from 14 day old plants grown from gemmae. The tissue was frozen in liquid nitrogen in Precellys tubes (Bertin Instruments, Montigny-le-Bretonneux, France) with 2.8mm Stainless steel beads (Bertin Corp., Rockville, MD, USA) and disrupted with a Precellys Evolution tissue homogenizer (Bertin Instruments) using the following settings: 7200RPM 10s, 5s pause, repeated thrice. RNA was extracted using a Spectrum Plant Total RNA kit (Sigma-Aldrich, Merck KGaA, Darmstadt, Germany).

Embryo samples were collected by hand dissection, with one embryo per replicate (fig. S1). Embryos and the surrounding maternal calyptra tissue were dissected out of the archegoniophore into 10% RNALater (Qiagen, Hilden, Germany) on Microscope slides with cavities (Marienfeld Superior, Lauda-Königshofen, Germany) and the embryo was further dissected out of the surrounding maternal tissue. Each embryo was washed four times in a series of wells containing 150μL 10% RNALater to remove any maternal RNAs, as previously described for the pure isolation of plant embryos (Kao & Nodine, 2020). Each embryo was then placed in 30μL 100% RNALater on ice until sample collection was completed. The solution was diluted to 10% RNALater by the addition of 270μL RNase-free water (Zymo Research, Irvine, CA), vortexed gently, and the solution removed. Samples were either resuspended in 30μL 100% RNALater and stored at -70°C or in 100μL TRI reagent (Zymo Research). Samples were crushed with a micropestle and RNA was extracted using a Direct-zol RNA MicroPrep kit (Zymo Research). All RNA samples were treated to remove DNA using a DNase Treatment and Removal kit (Invitrogen, Thermo Fisher Scientific Inc, Waltham, MA, USA).

RNA-seq libraries were generated from total RNA following the Smart-seq2 protocol (Picelli et al., 2014). cDNA synthesis was performed on 1μL of total RNA. 1μL of 10μM 5’-Bio-anchored oligo dT ([Btn]AAGCAGTGGTATCAACGCAGAGTACTTTTTTTTTTTTTTTTTTTTTTTTTTTTT TVN) and 1μL 10mM dNTPs were added to each sample and incubated at 72°C for 3 minutes and immediately placed on ice. 7μL of a mastermix containing 0.5μL SuperScript IV (Invitrogen), 0.25μL RiboLock (ThermoFisher Scientific), 2μL Superscript IV buffer (Invitrogen), 1μL 100mM DTT (Invitrogen), 2μL 5M Betaine (Sigma-Aldrich), 0.9μL MgCl_2_, 0.1μL nuclease-free water (Invitrogen), and 1μL 10μM 5’-Bio TSO ([Btn]AAGCAGTGGTATCAACGCAGAGTACrGrG+G, Exiqon) was added to each sample and the cDNA synthesis reaction took place under the following thermocycling conditions: 42°C 90min, <50°C 2min, 42°C 2min>x10, 70°C 15min. 40μL of a mastermix containing 14.5μL nuclease-free water, 25μL Q5 Hot Start 2x MasterMix (New England Biolabs, Ipswich, MA), and 0.5μL 10μM 5’ Bio-ISPCR oligo ([Btn]AAGCAGTGGTATCAACGCAGAGT) was added to each sample and the PCR pre-amplification took place under the following thermocycling conditions: 98°C 3min, <98°C 15s, 67°C 20s, 72°C 6min>x12, 72°C 5 minutes. Samples were cleaned up by bead purification using 1x volume of MBSPure beads (Molecular Biology Services, IMP, Vienna, Austria) and samples were eluted in 15μL of 10mM Tris-HCl. 5-50ng of each sample was used for the tagmentation reaction, containing 2.5μL of 4x TAPS-DMF buffer and 1μL of Tn5 (Molecular Biology Services, IMP), which was 3 minutes at 55°C, after which samples were immediately placed on ice. Samples were purified using a DNA Clean and Concentrator kit (Zymo Research) using manufacturer’s instructions and eluted in 10μL of 10mM Tris-HCl. Tagmented samples were amplified by the addition of 2.5μL each of 10μM barcoded forward and reverse primers (Picelli et al., 2014) and 15μL Q5 2x HiFI MasterMix (New England Biolabs) using the following thermocycling conditions: 72°C 3 minutes, 98°C 20s, <98°C 10s, 63°C 30s, 72°C 3 minutes>x5. Amplified samples were cleaned up by bead purification using 1x volume of MSBPure beads (Molecular Biology Services, IMP).

Samples were sequenced on an Illumnia NovaSeq to generate 50bp paired-end reads. Three biological replicates each of male (Tak-1) and female (Cam-2) vegetative tissue, 11 of wild-type (Cam-2 x Tak-1) embryos, and 17 of mutant (Cam-2 *e(z)2/e(z)3* x Tak-1) embryos were used for subsequent analyses.

### Chromatin profiling by CUT&RUN

Embryos and the surrounding calyptra were hand-dissected from archegoniophores and placed in Galbraith buffer (45mM MgCl_2_-6H2O, 30mM Trisodium citrate, 20mM MOPS) pH 7.0 plus 0.1% Triton X-100 and 1x cOmplete Protease Inhibitor Cocktail (Roche, Mannheim, Germany) on ice. Samples were crushed using a mortar and pestle on ice to release nuclei and were filtered through a 40µm filter (VWR, Radnor, PA, USA) before staining with 2µg/mL DAPI. Nuclei were sorted on a BD FACSARIA III (BD Biosciences, San Jose, CA, USA) to discriminate diploid embryonic nuclei from haploid maternal nuclei. Samples were sorted into 100µL of Wash buffer (20mM HEPES pH 7.5, 150mM NaCl, 0.5mM Spermidine, 1x cOmplete Protease Inhibitor Cocktail (Roche)) with 40,000 nuclei per replicate. Bio-Mag Plus Concanavalin A coated beads (Polysciences, Inc., Warrington, PA, USA) were activated by mixing 10µL per sample of ConA beads in 1.5mL Binding buffer (20mM HEPES-KOH pH 7.9, 10mM KCl, 1mM CaCl_2_, 1mM MnCl_2_). The beads were placed on a magnet, liquid was removed, and the beads were resuspended in 1.5mL Binding buffer. Liquid was again removed from the beads on a magnet and beads were resuspended in 10µL Binding buffer per sample. 10µL of the activated beads were added to each sorted nuclei sample and incubated at room temperature for 10min on a rotator. Liquid was removed from the bead-bound nuclei on a magnet and samples were resuspended in 50µL Antibody buffer (Wash buffer plus 2mM EDTA). 0.5µL of each antibody (H3K27me3 Millipore, Temecula, CA, USA, #07-449 RRID:AB_310624; H3K36me3 Abcam, Cambridge, UK, ab9050 RRID:AB_306966; H3K9me1 Abcam ab9045 RRID:AB_306963; H3 Abcam ab1791 RRID:AB_302613) used was added to samples while gently vortexing and samples were incubated overnight at 4°C on a shaker. Liquid was removed from the samples on a magnet and washed twice in 1mL Wash buffer before resuspending in 50µL Wash buffer. 1.16µL of 30µg/mL pAG-MNase (Molecular Biology Service, IMP) was added to each sample with gently vortexing and placed on a shaker for 10min at room temperature. Liquid was removed from the samples on a magnet and washed twice in 1mL Wash buffer before resuspending in 150µL Wash buffer. 3µL 100mM CaCl_2_ was added to ice-cold samples while gently vortexing and shaken at 4°C for two hours. 100µL STOP buffer (340mM NaCl, 20mM EDTA, 4mM EGTA, 50μg/mL RNase A (ThermoFisher Scientific), 50μg/mL glycogen, 10pg/mL heterologous HEK293 DNA) was added to stop the reaction. Samples were incubated at 37°C for 10min at 500RPM then spun at 4°C for 5min at 16,000G. Samples were placed on a magnet and the liquid containing released DNA fragment was transferred to a new tube. 2.5µL 10% SDS and 2.5µL 20mg/mL Proteinase K (ThermoFisher Scientific) was added to each sample, mixed by inversion, and incubated for 1hr at 50°C. 250µL buffered phenol-chloroform-isoamyl solution (25:24:1) was added to each sample, followed by vortexing and transfer to MaXtract tubes (Qiagen). Samples were spun for 5min at 16,000G. 250µL chloroform was added and samples were spun for 5min at 16,000G. The top aqueous phase was transferred to a fresh tube containing 2µL 2mg/mL glycogen. 625µL 100% EtOH was added before vortexing and chilling at -20°C overnight. DNA extraction continued with spinning for 10min at 4°C at 20,000G. The supernatant was poured off and 1mL 100% EtOH was added to the samples before spinning again for 1min at 4°C at 16,000G. Supernatant was discarded and samples air-dried before dissolving in 50µL 0.1x TE. A NEBNext Ultra II DNA library prep kit for Illumina (New England Biolabs) was used according to the manufacturer’s instructions for sample library preparation. Samples were sequenced on either and Illumina HiSeqv4 or NovaSeq to generate 50bp paired-end reads. Two biological replicates were used for each sample for H3K27me3, H3K36me3, H3K9me1 and H3 in wild-type (Cam-2 x Tak-1) embryos and mutant (Cam-2 *e(z)2/e(z)3* x Tak-1) embryos.

### Whole genome sequencing

Whole genome sequencing of Cam-2 was done as previously described (Iwasaki et al., 2021). 5g of 14 day old Cam-2 plants grown from gemmae were collected and frozen in liquid nitrogen. Samples were crushed using a mortar and pestle on ice and ground further in 25mL PVPP buffer (50mM Tris-HCl pH 9.5, 10mM EDTA, 4M NaCl, 1% CTAB, 0.5% PVPP, 1% beta-mercaptoethanol). The mixture was divided into two 50mL Falcon tubes and incubated at 80°C for 30 minutes in a water bath. Samples were cooled to room temperature and 7.5mL chloroform was added to each tube, followed by 5mL TE-saturated phenol after mixing. Samples were spun at 20,000G for 5 minutes at room temperature and the upper aqueous phase was transferred to a new 50mL tube. 1x volume of water and 4x volume of 100% EtOH were added and mixed, and samples were frozen at -70°C. Tubes were thawed and spun at 20,000G at 4°C for 15 minutes. The supernatant was poured off, samples were spun again briefly, and the remaining supernatant pipetted off. 2mL of 1x TE was added to each tube and incubated at 60°C for 10 minutes without mixing. The supernatant was transferred to another tube and incubated at 60°C for 10 minutes without mixing. 2μL of RNaseA (ThermoFisher Scientific) was added and samples incubated at 37°C for 5 minutes. 500μL was split into 2mL tubes and 50μL of 3M sodium acetate pH 5.2 and 1mL 100% EtOH were added. Samples were incubated at -20°C for 30min and spun at 13,000RPM for 15min. After removing the supernatant, pellets were rinsed twice with 1mL 70% EtOH and spun at 13,000RPM for 5min. Pellets were dried for 90s at 65°C and resuspended in 1mL 1xTE.

Library preparation was done by tagmentation. Briefly, 1µL gDNA was mixed with 2.5µL 4x TAPS-DMF buffer and 5µL activated Tn5 (Molecular Biology Services, IMP). Tagmentation proceeded for 5min at 55°C before cooling on ice. Samples were purified with a Zymo DNA Clean and Concentrator kit (Zymo Research) according to manufacturer instructions and eluted in 10µL 10mM Tris-HCl. PCR amplification was done by adding 2.5µL each of 10µM forward and reverse primers, plus 15µL NEBNext 2x HiFi PCR MasterMix (New England Biolabs) and thermocycling with the following conditions: 72°C 3min, 98°C 30s, <98°C 10s, 63°C 30s, 72°C 3min>x5. Samples were cleaned up by bead purification and sequenced on an Illumina NextSeq550 to generate 75bp paired-end reads.

### Interphase nuclei immunofluorescence slide preparation

Sporophytes were hand-dissected from archegoniophores and placed in Galbraith buffer (45mM MgCl2-6H2O, 30mM Trisodium citrate, 20mM MOPS) pH 7.0 plus 0.1% Triton X-100 and 1x cOmplete Protease Inhibitor Cocktail (Roche) on ice. Samples were crushed in a mortar and pestle on ice and filtered through a 40µm filter (VWR). 16% paraformaldehyde was added to reach a final concentration of 4% PFA and incubated on ice for 20min. Glycine was added to a final concentration of 125mM and nuclei were spotted onto glass slides and dried at room temperature for 20min.

### Mitotic cells immunofluorescence slide preparation

Sporophytes were hand-dissected from archegoniophores and placed in 1x PBS with 0.1% Triton-X 100 (PBST) and 4% paraformaldehyde on ice. Samples were fixed by applying a vacuum for 15min followed by 45min at 4°C. Samples were washed thrice for 10min each with PBST at 4°C with gentle shaking. Cell walls were digested by incubating samples in PBST plus 1% cellulase (Duchefa Biochemie) at 37°C for 10min in a damp chamber.

Samples were washed thrice for 10min each with PBST at 4°C with gentle shaking. Intact tissues were placed in 10µL PBST on a glass slide and squashed with a cover slip. Slides were dipped in liquid nitrogen and the cover slip was removed with a razor blade.

### Zygotic cells immunofluorescence slide preparation

To obtain archegonia holding synchronized zygote, fertilization timing was synchronized using an in vitro fertilization method described previously (Hisanaga et al., 2021). At 3 daf, archegoniophores were dissected under a Lynx EVO stereomicroscope (Vision Engineering, Woking, UK) and clusters of archegonia were collected into Fixative buffer (4% PFA, 1xPBS). Fixed tissues were then dehydrated and embedded in paraffin using the Donatello tissue processor (Diapath, Martinengo, Italy). Paraffin sectioning was done with a HM355S microtome (Microme, Walldorf, Germany) with 4μm thickness. Slides were deparaffinized and rehydrated with a Gemini autostainer (Fisher Scientific) with the following protocol: Xylene 5min, Xylene 5min, EtOH 100% 5min, EtOH 100% 5min, EtOH 95% 5min, EtOH 70% 5min, EtOH 30% 5min, Running tap water 1min, Water. Antigen retrieval was performed by boiling slides in Sodium Citrate buffer pH 6.0.

### Immunostaining of slides

Immunostaining of slides was done by an InsituPro VSi staining system (Intavis, Cologne, Germany) as previously described, with minor modifications (Borg, Buendía, & Berger, 2019). Slides were washed for 10min with TBS with 0.1% Tween-20 (TBST) 5 times then blocked with Blocking buffer (1x TBS, 0.1% Tween-20, 2% bovine serum albumin (BSA), 5% normal goat serum (NGS)) twice for 30min each. One antibody per slide (H3K27me3 Millipore #07-449 RRID:AB_310624; H3K36me3 Abcam ab9050 RRID:AB_306966; H3K9me1 Abcam ab9045 RRID:AB_306963) was diluted 1:100 and slides were incubated for 6hrs. After washing with TBST six times for 10min each, slides were incubated with a 1:500 dilution of secondary antibody (Goat Anti-Rabbit IgG H&L, Alexa Fluor® 488, ab150077 RRID:AB_2630356; Goat Anti-Mouse IgG H&L, Alexa Fluor® 568, ab175473 RRID:AB_2895153; Goat Anti-Rabbit IgG H&L, Alexa Fluor® 647, preadsorbed, ab150083 RRID:AB_2714032) and slides incubated for 2hrs. After eight 10min washes with TBST, slides were dried and counterstained with 1.5μg/mL 4′,6-diamidino-2-phenylindole (DAPI) and mounted in Vectashield antifade mounting medium with DAPI (Vector Laboratories, Piedmont, Italy) before being sealed with a coverslip and nail varnish.

### Immunofluorescence image acquisition

Images were acquired with a LSM 780 scanning laser confocal microscope (Zeiss).

### Combined Fluorescence in situ hybridization (FISH) and immunostaining method

Tissue fixation, nuclei isolation and flow cytometry were performed as described (N. Wang & Liu, 2020). A circular barrier was made with an ImmEdge™ Hydrophobic Barrier PAP Pen (Vector Laboratories) on the charged adhesion slide of a glass slide (ThermoFisher Scientific). Size of the circle was ∼ 0.7 cm diameter and ≥ 0.5 cm line thickness. Slides were dried for 30 min. 20μl of the nuclei suspension was transferred into PCR tubes and nuclei were incubated at 65°C for 30 min within a PCR Thermal Cycler. Heat shock treated nuclei were immediately transferred to ice for 5 min. 5 or 10μl of 0.1mg/ml RNase A (in 2x SSC buffer) was spotted into the circle drawn on the slide and mixed with 10µl (containing at least 1 x 10^4^ nuclei) of heat shock treated nuclei. The solution was spread within the circle barrier. Slides were incubated at 37°C in a ThermoBrite slide hybridizer (Leica Biosystems, Deer park, IL, USA) for 1 hour under a humid environment. At the end of this incubation, a very thin layer of solution remained on the glass slide. After incubation, the slides were treated for about 1min each by dipping up and down till the streaks go away in an ethanol series (100%, 95%, 90%, 80%, 60%, 30% EtOH). The slides were then treated in antigen retrieval buffer (10 mM sodium citrate pH 6.0) at room temperature for 5 min and then the antigen retrieval was started by boiling the slides for 10-12 min in a microwave at 700W. Slides were post-fixed in 4% formaldehyde solution 10 min after the slides cooled down to room temperature. After post-fixation, the slides were treated for about 1 min each by dipping up and down in ethanol series (30%, 60%, 80%, 90%, 95%, 100% EtOH). Slides were dried at room temperature for 1 hour.

The subsequent probe denaturation, hybridization, washing, and detection steps were performed according to (Bi et al., 2017) with minor changes. Anti-Histone H3 (mono methyl K9) antibody (Abcam, ab9045 RRID:AB_306963) or Anti-trimethyl-Histone H3 (Lys27) Antibody (Millipore, #07-449 RRID:AB_310624) was diluted 1:500 in antibody buffer (5% BSA, 4x SSC, 0.2% Tween 20). 10μl of the antibody mixture was pipetted onto the slides. The slides were incubated in a humid box at 37°C for 1 hour. After 1 hour of antibody binding, slides were washed for 5 min in a solution of 4x SSC with 0.2% Tween 20 in a foil-wrapped jar at room temperature on the shaker 3 times. 100μl 1:150 Anti-rabbit Alexa Fluor 546-conjugated goat antibody (Invitrogen, AB_2534093 RRID:AB_2534093) was dropped onto the slides. The slides were incubated at 37°C for 1 hour followed by 3 times (5 mins each) washing steps. Then, the slides were mounted with 5μl SlowFade™ Diamond Antifade Mountant (Invitrogen). Slides were covered with a coverslip and sealed with nail polish. Images were acquired with a LSM 710 scanning laser confocal microscope (Zeiss, Oberkochen, Germany).

### Probe labeling for FISH

Probes were labeled according to the Nick Translation-based DNA Probe Labeling method (Roche). Tak-1 and Tak-2 genomic DNA was extracted by CTAB method (Murray & Thompson, 1980). For U chromosome probe and U chromosome competition probe, the Tak-2 gDNA was used as template. For V chromosome probe and V chromosome competition probe the Tak-1 gDNA was used as template. Fluoroprobe labelling mix: For V/U chromosome probe (dATP, dCTP, dGTP, dTTP, Dig-dUTP); For V/U chromosome competition probe (dATP, dCTP, dGTP, dTTP). For U chromosome FISH, U-chromosome probe and 5 times V-chromosome competition probe were loaded. For V chromosome FISH, V-chromosome probe and 5 times U-chromosome competition probe were loaded.

### Tissue clearing and DAPI staining of zygotes

Tissue clearing and DAPI staining for 3 daf zygotes were done as described previously (Hisanaga et al., 2021). Stained samples were mounted in Vectashield antifade mounting medium with DAPI (Vector Laboratories). Images were taken by with a LSM780 confocal microscope (Zeiss).

### Nuclear envelope visualization

To observe the nuclear envelope of 3 daf zygotes, ECpro:SUN-GFP (Hisanaga et al., 2021) females were fertilized with wild-type sperm and zygotes were excised under a Lynx EVO stereomicroscope (Vision Engineering) and mounted in half-strength Gamborg B5 media without vitamins (Duchefa Biochemie) liquid medium. Samples were observed under a Nikon C2 confocal laser-scanning microscope (Nikon Instech, Tokyo, Japan).

### Mutant fitness analyses

Four gemmae from Cam-2 and Cam-2 *e(z)2/e(z)3* plants were grown together. Images of each gemmaling was taken at four, seven and ten days after planting using a Lynx EVO stereomicroscope (Vision Engineering). The area of each gemmaling was calculated using FIJI v2.0.0 (Schindelin et al., 2012) and plotted as a smoothed curve using the loess function and formula y∼x in R v3.5.1 (R Core Team, 2018) with the ggplot2 v3.3.5 package (Wickham, 2016).

Gemmae from Cam-2 and Cam-2 *e(z)2/e(z)3* were planted on Grodan and monitored until the first archegoniophores were visible. Pictures were taken after all replicates had produced archegoniophores to illustrate the synchronicity of archegoniophore developmental stage.

Images of fully dissected embryos (see Expression analysis sample collection section above for details) were taken with Lynx EVO stereomicroscope (Vision Engineering). The height and width of each embryo was calculated in FIJI v2.0.0 (Schindelin et al., 2012) using images of a calibration slide as reference. The sample area was calculated by multiplying height and width.

Aborted embryos can be identified by a browning of tissue, collapse of tissue within the calyptra, and the outgrowth of the perianth without growth of the embryo within. Embryo survival was calculated as the number of green, non-collapsed embryos per archegoniophore divided by the number of perianths with or without live embryos.

Mature embryos can be identified by the yellowing of tissue due to the production of spores within. The percentage of embryos producing spores was calculated as the number of mature yellow embryos per archegoniophore divided by the number of perianths with or without live embryos.

Spore germination was assessed by counting the number of sporelings growing out from spots of serially diluted spore solutions from single sporophytes. Mature embryos were dissected from archegoniophores, dried for one week, and frozen at -70°C. Frozen embryos were thawed and ruptured in 100µL sterile water using a sterile pipette tip. 80µL of the spore suspension was transferred to a tube containing 420µL sterile water. 500µL of 0.1% NaDCC (Sigma-Aldrich) was added to each sample and tubes were inverted and spun at 13,000RPM for 1 minute. The supernatant was removed, and spores were resuspended in 100µL sterile water. 20µL of spore suspension was spotted onto plates of half-strength Gamborg B5 media without vitamins (Duchefa Biochemie) and 1% agar. 20µL of spore suspension was carried to a tube containing 60µL sterile water. The process was repeated until dilutions of 1:1, 1:4, 1:16 and 1:64 were spotted. Images of sporeling germination and growth were taken at 11 days after planting.

### Transcriptome analysis

Published transcriptomes from male and female reproductive tissues, antheridiophores and archegoniophores, respectively (Higo et al., 2016) and wild-type Tak-2 x Tak-1 embryos (Frank & Scanlon, 2015) were downloaded from the SRA database.

Reads were mapped to the Takv6 genome (Iwasaki et al., 2021) wherein all SNP positions between Tak-1 and Cam-2 or between Tak-1 and Tak-2 were replaced with N’s, depending on the genotype of the sample (refer to SNP analysis section below). Reads were preprocessed with SAMtools v1.9 (H. Li et al., 2009) and BEDTools v2.27.1 (Quinlan & Hall, 2010), trimmed with Trim Galore (https://github.com/FelixKrueger/TrimGalore) and mapped with STAR v2.7.1 (Dobin et al., 2013). Transcripts per Million (TPM) values were calculated by RSEM v1.3.2 (B. Li & Dewey, 2011). Data from RSEM were imported into R v3.5.1 (R Core Team, 2018) using the tximport package v1.10.1 (Soneson, Love, & Robinson, 2015). Differential gene analysis was performed using DeSeq2 v1.22.2 (Love, Huber, & Anders, 2014). Principal component analysis was performed in R v3.5.1 (R Core Team, 2018). Effect size (Cohen’s *d*) was calculated in R using effsize v0.7.6 (Torchiano, 2020) where |d| < 0.2 is no effect, 0.2 < |d| < 0.5 is a small effect, 0.5 < |d| < 0.8 is a medium effect, and |d| > 0.8 is a large effect, as previously reported (Cohen, 1992). Heatmaps were generated in R using the pheatmap v1.0.12 package (Kolde, 2019).

### CUT&RUN data analysis

Reads were mapped to the Takv6 genome (Iwasaki et al., 2021) wherein all SNP positions between Tak-1 and Cam-2 were replaced with N’s (refer to SNP analysis section below). File processing and mapping parameters were performed as previously published (Montgomery et al., 2020). Chromatin enrichment per gene was calculated by counting the number of reads and normalizing to 1x genome coverage.

### SNP data analysis

Reads were preprocessed with SAMtools v1.9 (H. Li et al., 2009), BEDTools v2.27.1 (Quinlan & Hall, 2010) and Picard v2.18.27 (http://broadinstitute.github.io/picard/) before mapping to the Tak-1 genome with bwa v0.7.17 (H. Li & Durbin, 2009). SNPs were called using gatk v4.0.1.2 and the reference genome with all SNPs replaced with N’s was created (McKenna et al., 2010).

Mapped reads from CUT&RUN and RNA-Seq experiments were assigned to paternal or maternal genomes using SNPSplit v0.3.4 (Krueger & Andrews, 2016). Counts for the number of reads originating from either genome were calculated per sample using SAMtools v1.9(H. Li et al., 2009) and BEDTools v2.27.1 (Quinlan & Hall, 2010). The maternal ratio was determined by dividing the number of maternal reads by total reads per gene. For CUT&RUN data, only data from genes with more than ten total reads in each replicate were retained. For RNA-Seq data, only data from genes with more than fifty reads in total across all replicates were retained. Additionally, data from genes that were completely maternally biased in male Tak-1 RNA-Seq data or were completely paternally biased in female Cam-2 RNA-Seq data were excluded from further maternal ratio analyses.

### Interphase nuclei image deconvolution

Immunofluorescence images of interphase nuclei were deconvolved with Huygens Professional v21.04 (Scientific Volume Imaging B.V., Hilversum, The Netherlands) using the CMLE algorithm with 40 iterations and SNR values as follows: 6 for WT H3K27me3 samples H3K27me3 channel, 8 for WT H3K27me3 samples DAPI channel, 2 for WT H3K9me1 samples H3K9me1 channel, 4 for WT H3K9me1 samples DAPI channel, 4 for WT H3K36me3 samples H3K36me3 channel, 5 for WT H3K36me3 samples DAPI channel, 2 or 4 for mutant H3K27me3 samples H3K27me3 channel, 3 for mutant H3K27me3 samples DAPI channel.

### Interphase nuclei image nuclei segmentation

Nuclei were identified from DAPI signal marking DNA in each immunofluorescence image. An adaptive thresholding technique was used, based on the creation of a sequence of 20 threshold values spanning a range from a clearly too low threshold to a clearly too high threshold. A sequence of masks was thus obtained for each 3-dimensional image by thresholding it using these values. Subsequently, a maximum intensity projection of each mask was computed and size of each mask projection was evaluated. Typically, starting from the lowest threshold, such sequence first decreased rapidly, followed by a wide plateau, and ending by a decreasing tail near the highest threshold value (Figure 7A). The nearly constant plateau was detected by thresholding absolute value of neighbor size differences. Again, a sequence of thresholds was used, starting from a minimum equal to 1/10 of average of the differences, until the length of such plateau was larger than 6. Finally, the segmentation in the middle of the plateau was taken as the final one. In the last step eventual holes in the 3-dimensional mask were filled by a hole-filling operation and eventual thin gaps in the mask were filled by binary closing.

**Figure 7.**
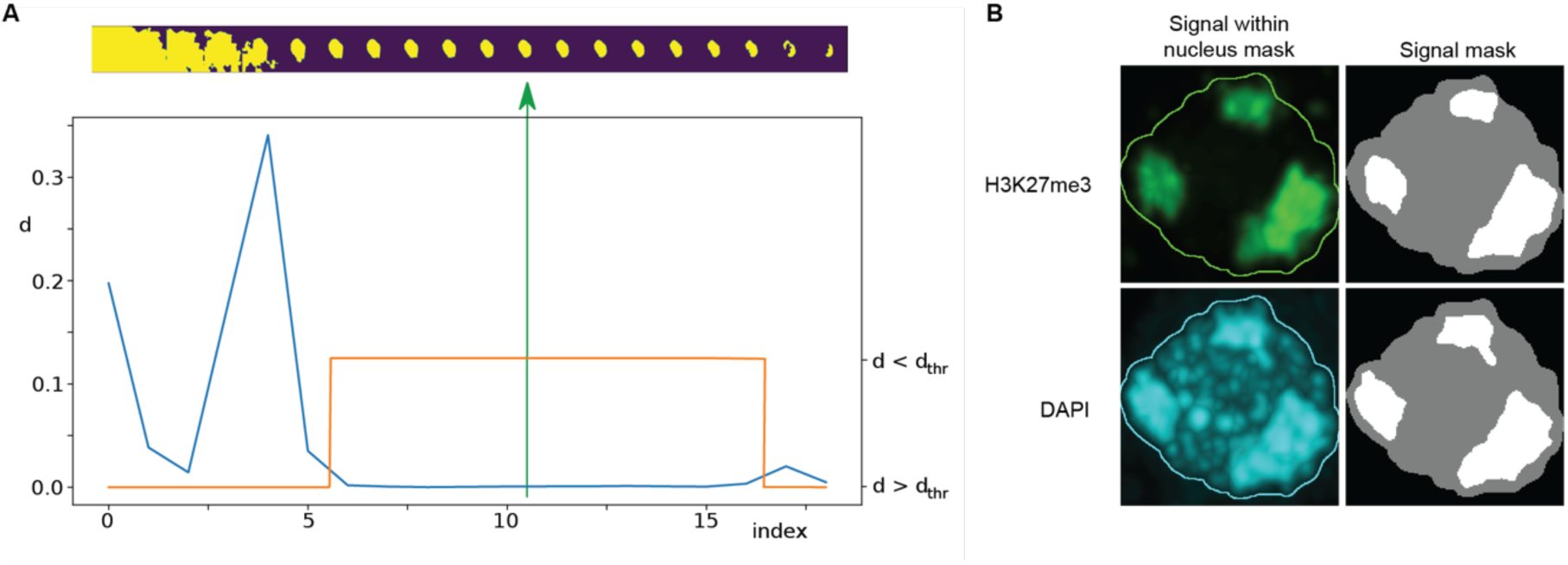
Immunofluorescence image analysis. (**A**) Nucleus segmentation. Top panel: A sequence of 20 segmentations created by thresholding. Bottom panel: Differences (d) between nucleus mask sizes (blue curve). Plateau identified by estimating a threshold value d_thr_ (orange curve). The selected segmentation is in the center of the plateau (green arrow).) Foci segmentation. Left: H3K27me3 and DAPI image bands with the overlayed nucleus border detected from the DAPI band. Right: Masks of the nucleus and detected foci.

Nucleus masks thus obtained were subsequently visually inspected. Except for a few cases, the nuclei were labeled correctly. The incorrect ones were manually adjusted by means of drawing functions in FIJI v2.0.0 (Schindelin et al., 2012).

### Interphase nuclei image foci segmentation

Foci within the nucleus area were detected separately for the DAPI and the immunofluorescence channels by k-means classification in 2, 3 and 4 classes, which in our case of single-valued data specified one, two or three threshold values. The foci mask was then computed by thresholding the input data using the highest of these thresholds. The results were visually inspected and classification in 3 classes was then taken for further processing and evaluation. Only the largest foci with size bigger than 20% of the nucleus size were considered (Figure 7B).

In a limited number of cases, one or two foci were missing, or foci were too large. In the first case there were two possibilities how to identify more foci: either by decreasing the expected foci size or by decreasing the threshold value. Thus, in a loop we multiplied both these values by a coefficient until the desired number of foci was reached. In the second case with too large foci, classification in 4 classes was used, which increased the highest threshold and simultaneously decreased the size of the foci.

### Statistical analyses

Statistical comparisons means of FISH immunostaining images and mutant growth analyses were performed with Wilcoxon tests in R v3.5.1 (R Core Team, 2018) with the ggpubr v0.4.0 package (Kassambara, 2020). Spearman correlations were calculated in R and with the ggpubr package.

## Acknowledgements

We thank A. Pauli, A. Burga, and I. Patten for suggestions and critical reading of the manuscript. F.B. acknowledges support from the PlantS, next-generation sequencing and histopathology facilities at the Vienna BioCenter Core Facilities (VBCF), and the BioOptics facility and Molecular Biology Services from the Institute for Molecular Pathology (IMP). This work was funded by FWF grants P26887, P28320, P32054, P33380 to F.B., FWF doctoral school DK W1238 to S.A.M., and European Research Council under the European Union’s Horizon 2020 research and innovation programme 757600 to C.L.

## Author contributions

S.A.M. and F.B. conceived and designed the experiments. S.A.M. performed the whole genome sequencing, RNA-seq, CUT&RUN, embryonic nuclei immunostaining, and fitness measurements. T.H. performed zygotic nuclei immunostaining. T.H. and S.A. generated material used in this study. N.W. performed immuno-FISH experiments. E.A. and S.A.M. performed statistical analyses and curated data. M.S. and S.A.M. analyzed image data. F.B. and C.L. supervised the study. S.A.M. and F.B. wrote the manuscript with input from C.L. and T.H.

## Data and materials availability

The CUT&RUN and RNA-seq sequencing datasets generated for the current study will be made available in the Gene Expression Omnibus (GEO) upon publication. Whole-genome sequencing data are deposited under BioProject accession number PRJNA795113. Publicly available datasets can be accessed under the DDBJ Sequence Read Archive accession numbers DRR050346-DRR050348 and DRR050351-DRR050353 and the NCBI Sequence Read Archive accession numbers SRR1553297-SRR1553299. Source data are provided with this paper. Original images are deposited online at FigShare and are publicly available as of the date of publication. DOI: 10.6084/m9.figshare.19249622 and 10.6084/m9.figshare.19249643. All original code has been deposited online at FigShare and is publicly available as of the date of publication. DOI: 10.6084/m9.figshare.19249592.

Any additional information required to reanalyze the data reported in this paper is available from the lead contact upon request. Further information and requests for resources and reagents should be directed to and will be fulfilled by the lead contact, Frédéric Berger.

## Supplementary Files

### Supplementary Figures

**Figure 1-figure supplement 1.**
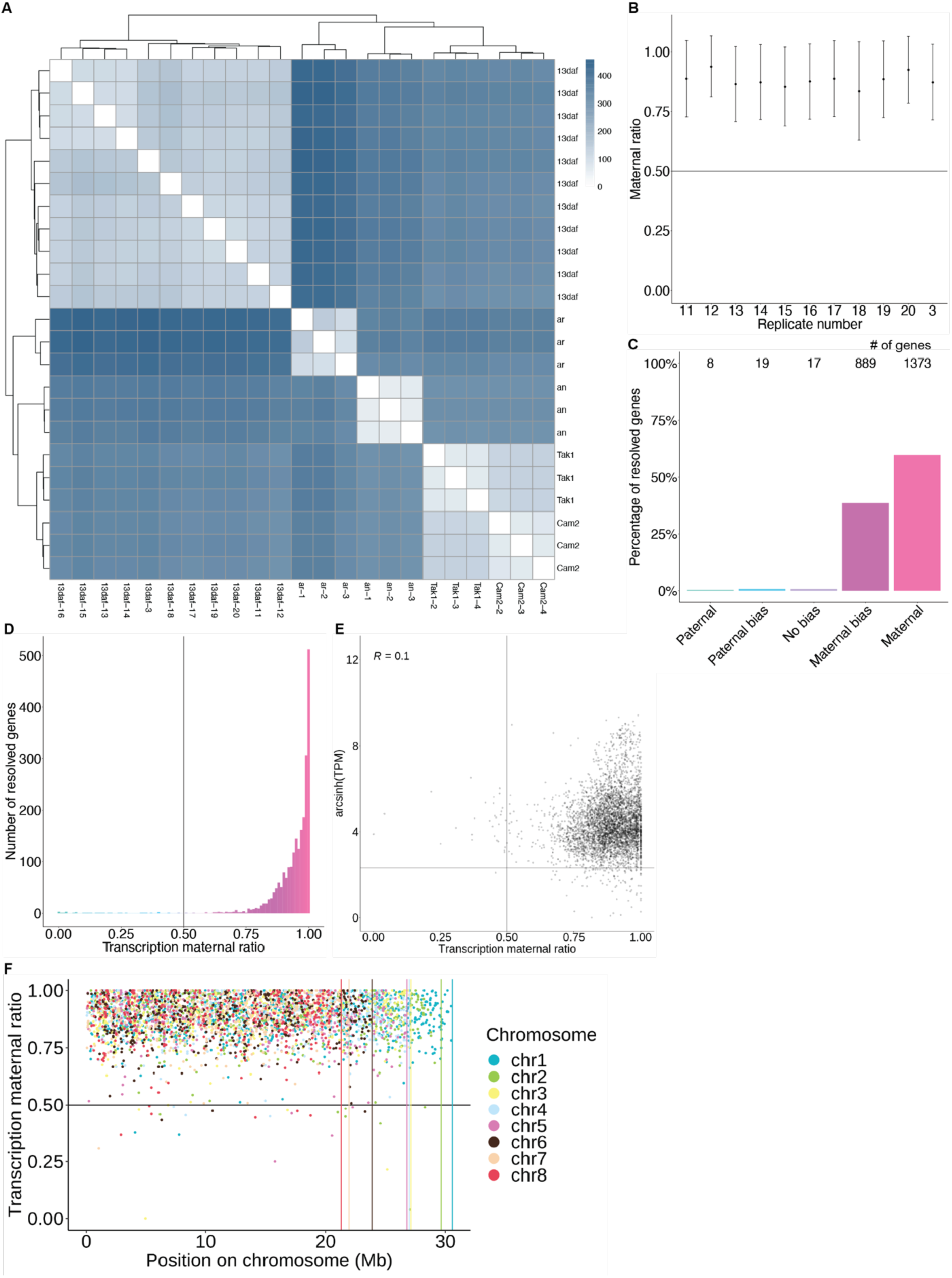
Maternally biased transcription in embryos. (**A**) Distance matrix heatmap of RNA-seq replicates. Individual replicates indicated along the bottom axis and the sample stage is indicated along the right axis. daf (days after fertilization), an (antheridiophore, male sexual organ), ar (archegoniophore, female sexual organ. Hierarchical clustering of replicates indicated along top and left axes. Computed distance indicated by scale bar. (**B**) Maternal ratio of embryo RNA-seq replicates. Black dots indicate the mean maternal ratio of transcription over all resolved genes per replicate. Black vertical lines indicate ± standard deviation. (**C**) Percentage of measured genes within each category of maternal ratio (*p_m_*) of transcription in wild-type (Tak-2 x Tak-1) embryos. Segments are for paternal (*p_m_* < 0.05), paternal bias (0.05 < *p_m_* ≤ 0.35), no bias (0.35 < *p_m_* < 0.65), maternal bias (0.65 ≤ *p_m_* < 0.95), and maternal (0.95 ≤ *p_m_*) expression of genes, with the number of genes indicated above each bar. (**D**) Histogram of the maternal ratio (*p_m_*) of transcription per gene in wild-type (Tak-2 X Tak-1) embryos. Each bin is 0.01 units wide. (**E**) Scatterplot of gene expression (arcsinh transformed Transcripts per Million (TPM)) versus transcription maternal ratio per gene in wild-type (Cam-2 X Tak-1) embryos. Spearman correlation is indicated. (**F**) Transcription maternal ratio per gene along the length of each chromosome. Vertical lines indicate the end of each chromosome.

**Figure 2-figure supplement 1.**
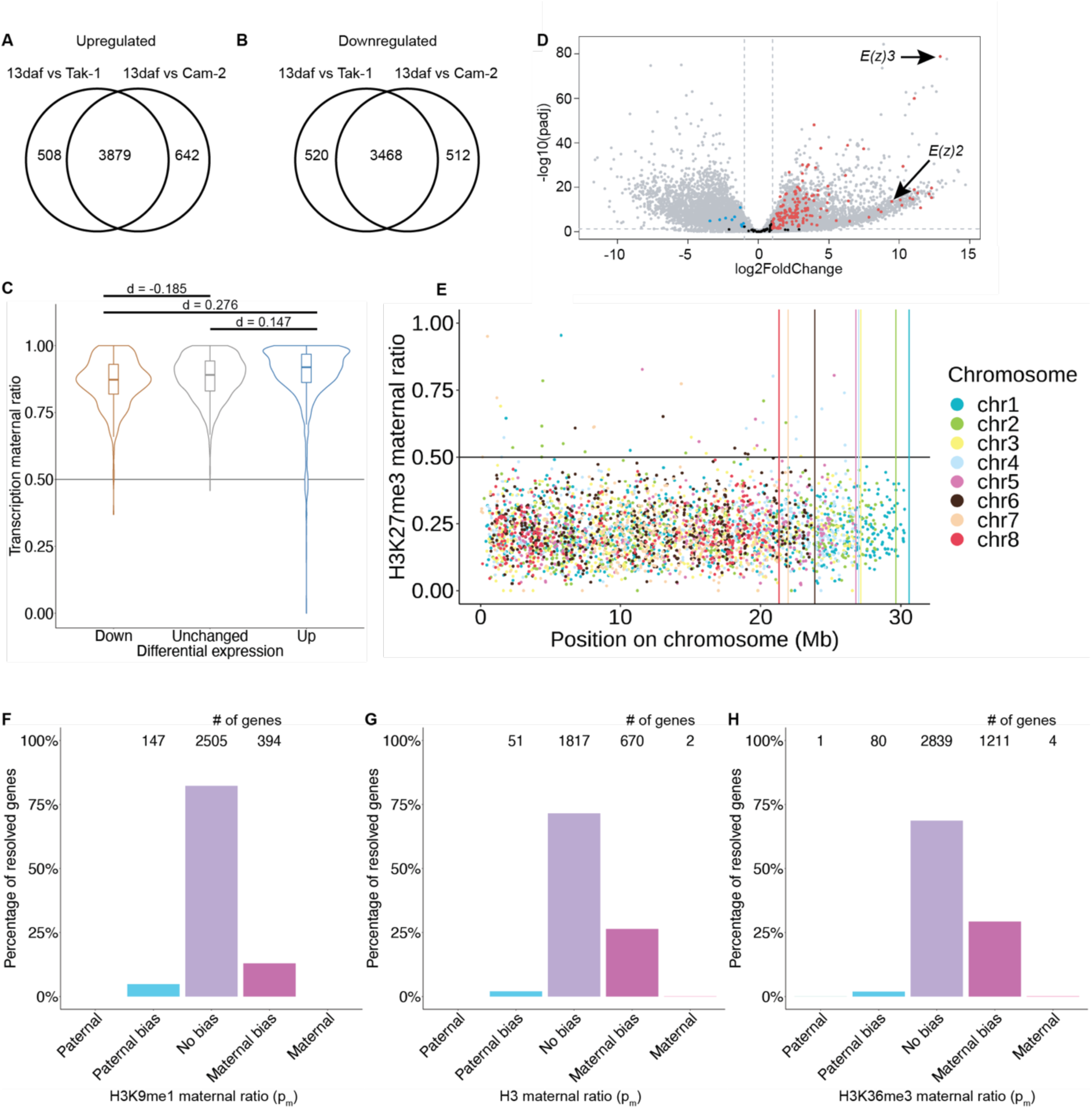
Paternally biased H3K27me3 in embryos. (**A**) Venn diagram of upregulated genes in wild-type embryos compared to male (Tak-1) and female (Cam-2) parents. (**B**) Venn diagram of downregulated genes in wild-type embryos compared to male (Tak-1) and female (Cam-2) parents. (**C**) Violin plots of maternal ratio of transcription for shared differentially expressed genes in embryos versus wild-type vegetative tissue from parents. Cohen’s *d* effect size values are indicated for pairwise comparisons of Down to None, Up to None, and Up to Down where |d| < 0.2 is no effect and 0.2 < |d| < 0.5 is a small effect, as previously reported (Cohen, 1992). (**D**) Volcano plot of a differential gene expression analysis between wild-type embryos and the male parent. The negative log transformed p-value per gene is plotted against the log2 fold-change in expression. Vertical dashed lines indicate a log2 fold-change of -1 and 1. The horizontal dashed line indicates a p-value of 0.05. Dots not in grey indicate chromatin-related genes, blue for significantly downregulated genes, red for significantly upregulated genes, black for genes not significantly downregulated nor upregulated. (**E**) H3K27me3 maternal ratio per gene along the length of each chromosome. Vertical lines indicate the end of each chromosome. (**F**-**H**) Percentage of measured genes within each category of maternal ratio (*p_m_*) of (**F**) H3K9me1 in wild-type embryos. (**G**) H3, and (**H**) H3K36me3 in wild-type embryos. Segments are for full paternal (*p_m_* < 0.05), paternal bias (0.05 < *p_m_* ≤ 0.35), no bias (0.35 < *p_m_* < 0.65), maternal bias (0.65 ≤ *p_m_* < 0.95), and full maternal (0.95 ≤ *p_m_*) chromatin enrichment of genes, with the number of genes indicated above each bar.

**Figure 3-figure supplement 1.**
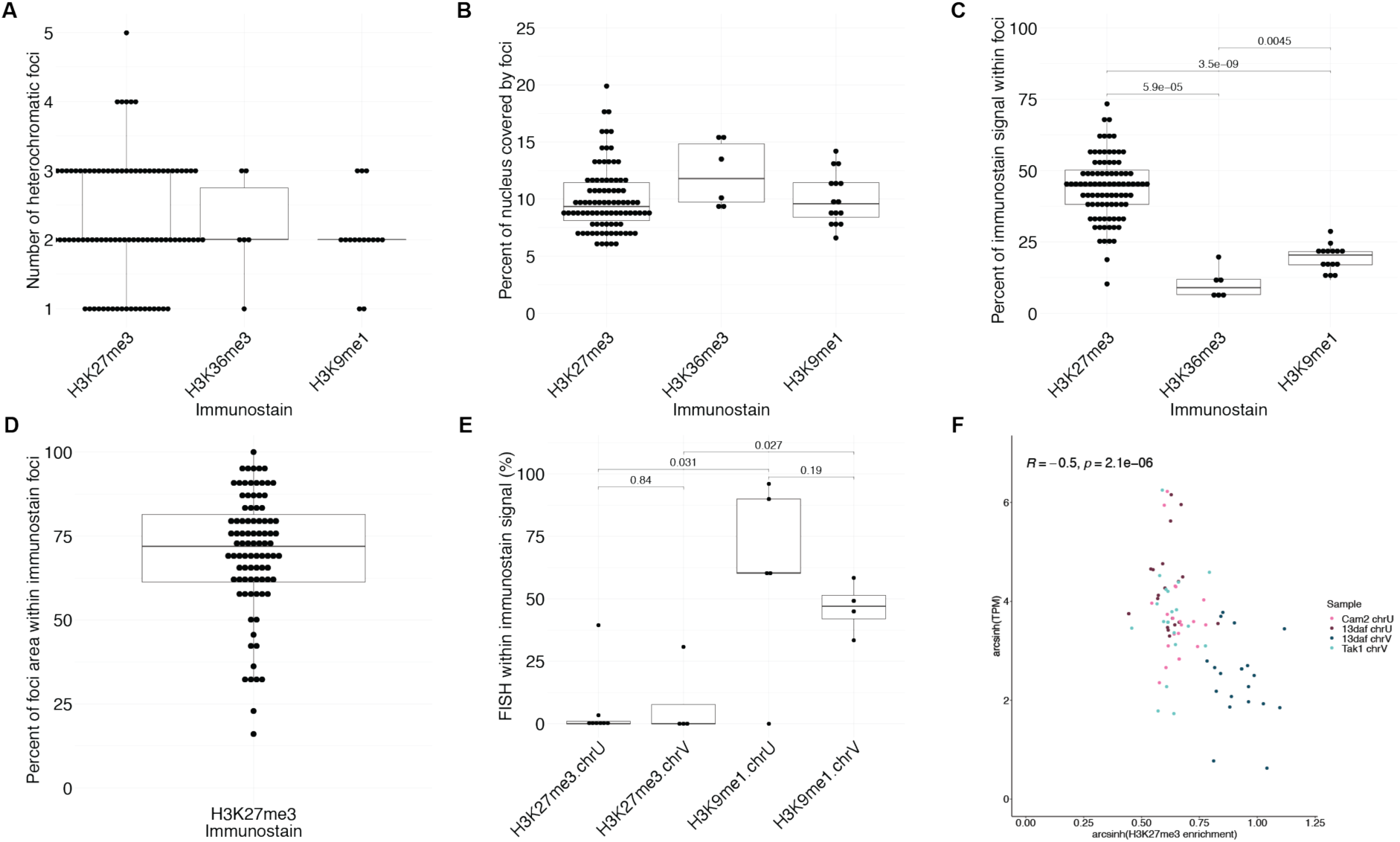
Quantification of immunofluorescence experiments. (**A**) Number of heterochromatic foci per nucleus in wild-type embryos. (**B**) Percentage of nuclear area covered by heterochromatic foci in wild-type embryos. (**C**) Percentage of immunofluorescence signal located within heterochromatic foci in wild-type embryos. *P* values are indicated, unpaired two-tailed Wilcoxon test. (**D**) Percentage of heterochromatic foci area overlapping with H3K27me3 foci in wild-type embryos. (**E**) Quantification of sex chromosome FISH signal located within H3K27me3 or H3K9me1 heterochromatic foci. *P* values are indicated, unpaired two-tailed Wilcoxon test. (**F**) Scatterplot of arcsinh transformed TPM values versus arcsinh transformed H3K27me3 enrichment for sex chromosome gametologs in vegetative and embryonic samples. Spearman correlation and *P* value are indicated.

**Figure 4-figure supplement 1.**
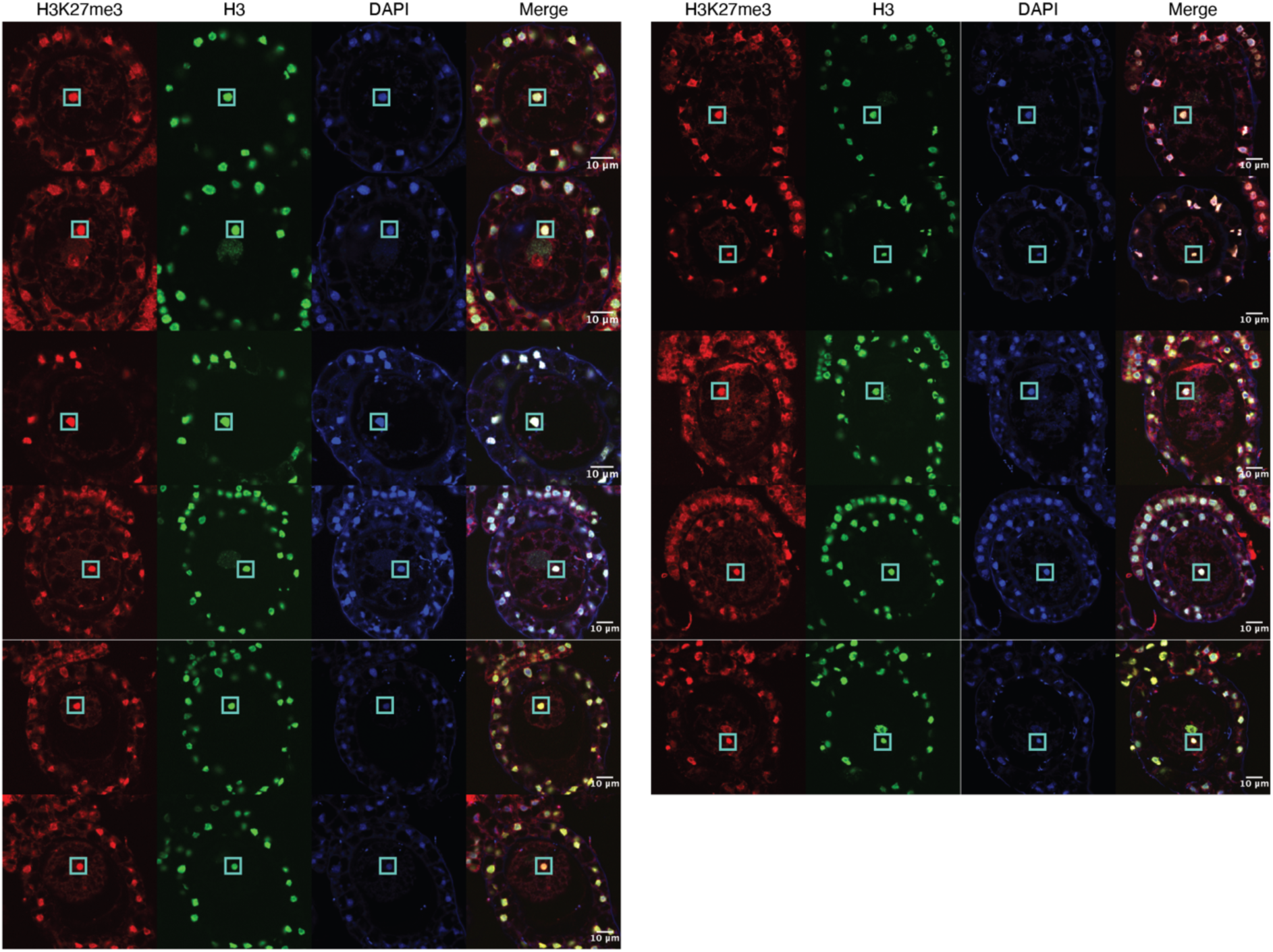
Immunofluorescence of pronuclei. Images of wild-type zygotes at 3 days after fertilization immunostained against H3K27me3 (red) and H3 (green) and counterstained with DAPI (blue). Paternal pronuclei are indicated with a cyan box in each image. Contrast is enhanced for each image and channel independently for visualization purposes. Scale bars as indicated.

**Figure 5-figure supplement 1.**
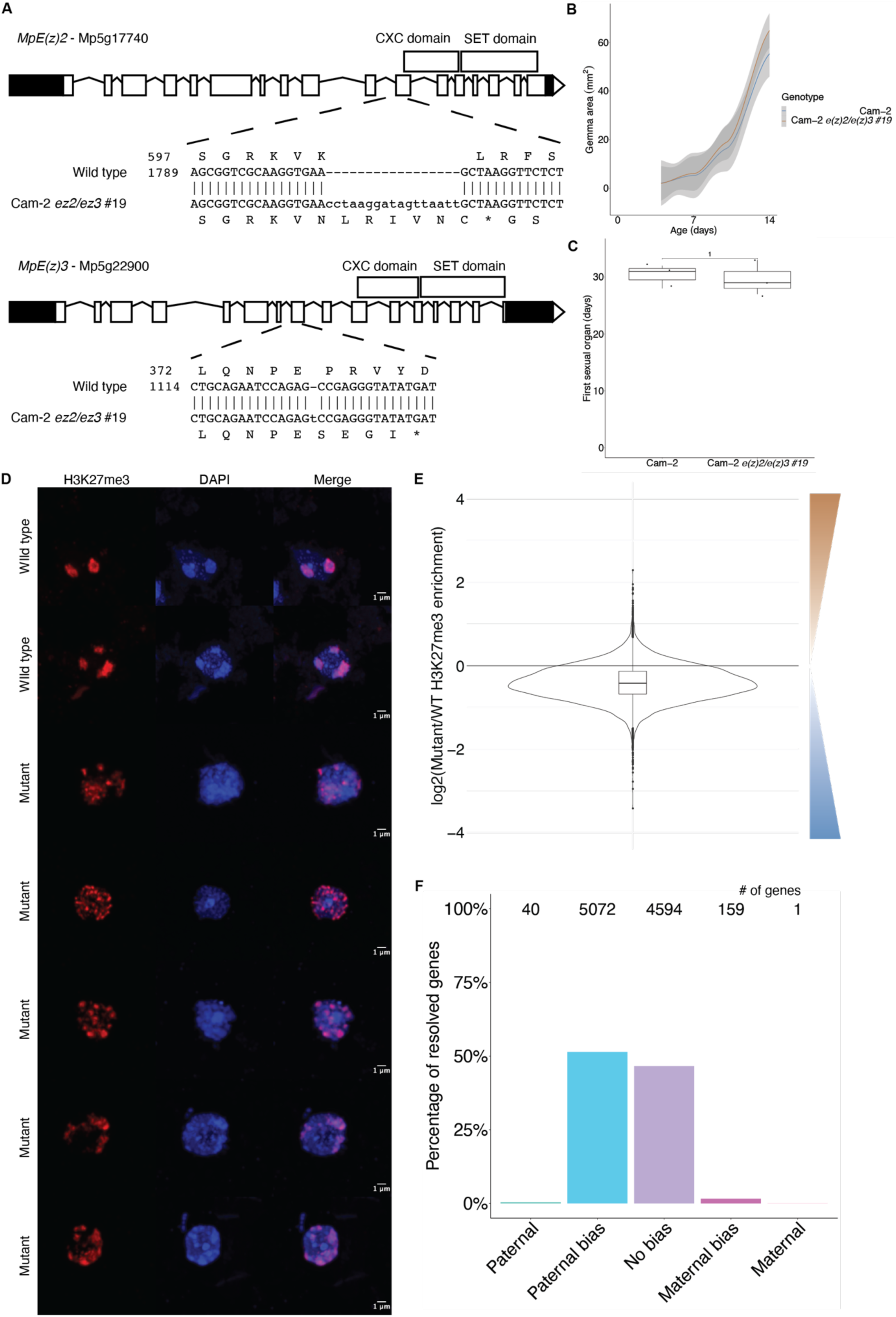
Chromatin phenotypes of e(z)2/e(z)3 mutants. (**A**) Structure of *E(z)2* and *E(z)3* genes and the mutations generated in each. (**B**) Growth of Cam-2 *e(z)2/e(z)3* mutants in the vegetative stage relative to Cam-2 wild type. Area of vegetative haploid plant growth as a function of the number of days after planting gemmae (propagules). (**C**) Timing of the appearance of sexual organs in Cam-2 *e(z)2/e(z)3* mutants relative to Cam-2 wild type as a function of the number of days after planting. *P* value is indicated, unpaired two-tailed Wilcoxon test. (**D**) Representative set of wild-type and mutant embryo immunofluorescence images. Images are maximum intensity projections. Scale bars are as indicated. (**E**) Log2 ratio of H3K27me3 enrichment between mutant and wild type. Brown scale indicates greater H3K27me3 enrichment in the mutant, whereas the blue scale indicates greater H3K27me3 enrichment in wild type. (**F**) Percentage of measured genes within each category of maternal ratio (*p_m_*) of H3K27me3 in mutant embryos. Segments are for full paternal (*p_m_* < 0.05), paternal bias (0.05 < *p_m_* ≤ 0.35), no bias (0.35 < *p_m_* < 0.65), maternal bias (0.65 ≤ *p_m_* < 0.95), and full maternal (0.95 ≤ *p_m_*) H3K27me3 of genes, with the number of genes indicated above each bar.

**Figure 5-figure supplement 2.**
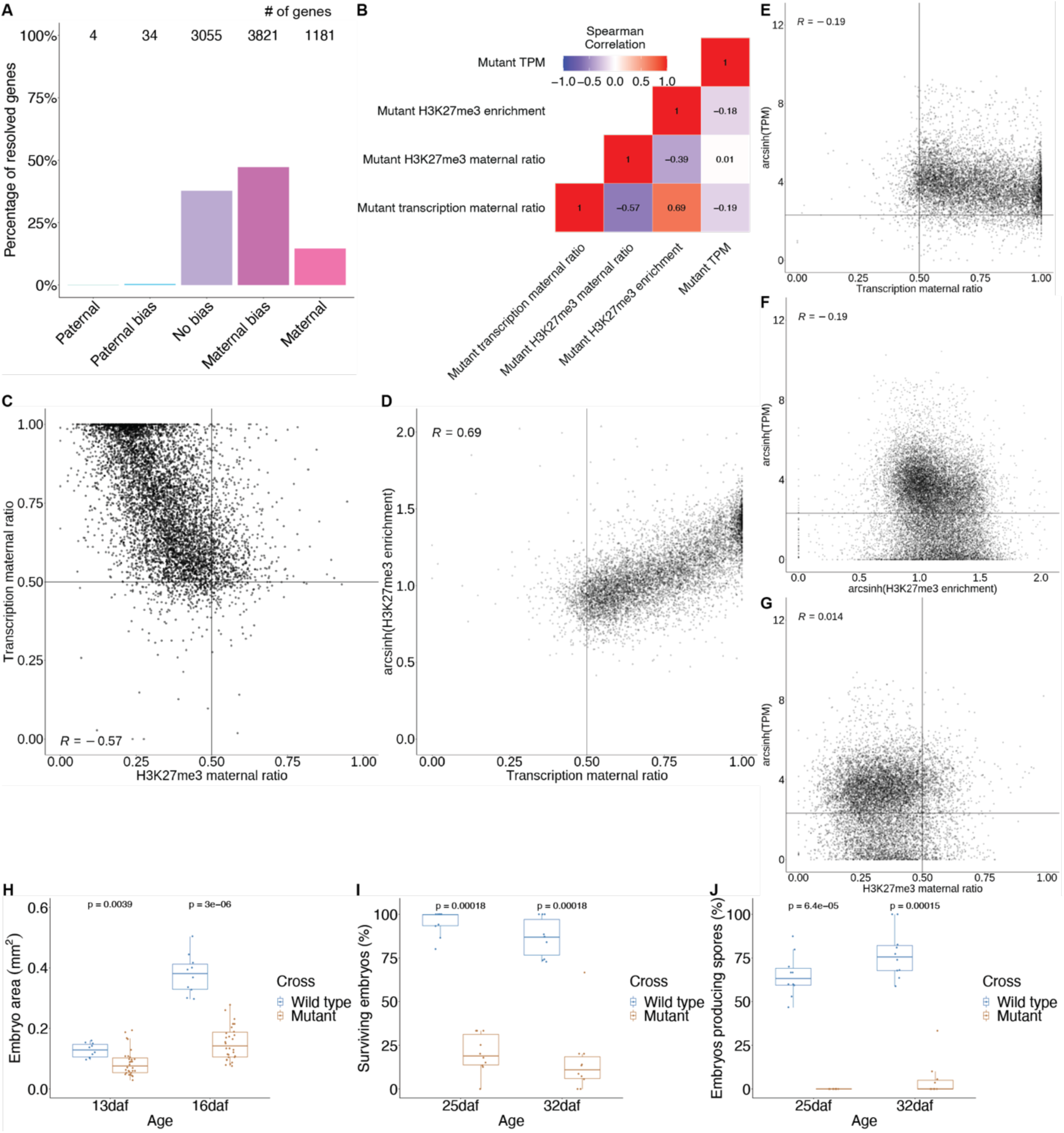
Transcription phenotypes in mutant embryos. (**A**) Percentage of measured genes within each category of maternal ratio (*p_m_*) of transcription in mutant embryos. Segments are for full paternal (*p_m_* < 0.05), paternal bias (0.05 < *p_m_* ≤ 0.35), no bias (0.35 < *p_m_* < 0.65), maternal bias (0.65 ≤ *p_m_* < 0.95), and full maternal (0.95 ≤ *p_m_*) expression of genes, with the number of genes indicated above each bar. (**B**) Heatmap of Spearman correlations of gene features in mutant embryos. (**C**) Scatterplot of transcription maternal ratio versus H3K27me3 maternal ratio per gene in mutant embryos. (**D**) Scatterplot of arcsinh transformed H3K27me3 enrichment versus transcription maternal ratio per gene in mutant embryos. (**E**) Scatterplot of arcsinh transformed Transcript per Million (TPM) values versus transcription maternal ratio per gene in mutant embryos. (**F**) Scatterplot of arcsinh transformed TPM values versus arcsinh transformed H3K27me3 enrichment per gene in mutant embryos. (**G**) Scatterplot of arcsinh transformed TPM values versus H3K27me3 maternal ratio per gene in mutant embryos. Spearman correlations are indicated for each scatterplot. (**H**) Embryo size of wild-type and mutant embryos 13 and 16 days after fertilization (daf) measured by the area of a bounding box. (**I**) Percentage of wild-type and mutant embryos per female sex organ that survive to maturity at 25 and 32 daf. (**J**) Percentage of wild-type and mutant embryos per female sex organ that have produced spores at 25 and 32 daf. *P* values are indicated, unpaired two-tailed Wilcoxon test.

### Supplementary data

Supplemental Video 1: Movie of the dissection of a representative Marchantia embryo from surrounding calyptra of maternal origin.

Supplemental Table 1: List of Marchantia chromatin-related genes and their expression status in embryos relative to other tissues.

